# A derivation error that affects carbon balance models exists in the current implementation of the modified Arrhenius function

**DOI:** 10.1101/2020.01.27.921973

**Authors:** Bridget Murphy, Joseph R. Stinziano

## Abstract

- Understanding biological temperature responses is crucial to predicting global carbon fluxes. The current approach to modelling temperature responses of photosynthetic capacity in large scale modelling efforts uses a modified Arrhenius equation.
- We rederived the modified Arrhenius equation from the source publication from 1942 and uncovered a missing term that was dropped by 2002. We compare fitted temperature response parameters between the correct and incorrect derivation of the modified Arrhenius equation.
- We find that most parameters are minimally affected, though activation energy is impacted quite substantially. We then scaled the impact of these small errors to whole plant carbon balance and found that the impact of the rederivation of the modified Arrhenius equation on modelled daily carbon gain causes a meaningful deviation of ~18% day^−1^.
- This suggests that the error in the derivation of the modified Arrhenius equation has impacted the accuracy of predictions of carbon fluxes at larger scales since >40% of Earth System Models contain the erroneous derivation. We recommend that the derivation error be corrected in modelling efforts moving forward.

## Introduction

Globally, photosynthesis and autotrophic respiration are the largest biological carbon fluxes, with photosynthesis removing ~120 Gt C year^−1^ from the atmosphere and autotrophic respiration releasing ~60 Gt C year^−1^ back to the atmosphere (Amthor, 2000; Ciais *et al.*, 2013). Given the temperature sensitivity of these large carbon fluxes (Lombardozzi *et al.*, 2015), understanding how photosynthesis and respiration respond on acute, acclimatory, and adaptive timescales is crucial for predicting vegetative and carbon cycle responses to future global climates (Rogers *et al.*, 2017; Stinziano *et al.*, 2018). Biological temperature responses including photosynthesis and respiration are typically assumed to be exponential or peaked (Way & Yamori, 2014; Smith & Dukes, 2017; Kumarathunge *et al.*, 2019). Exponential responses are usually modelled based on an Arrhenius-type curve (Arrhenius, 1915):

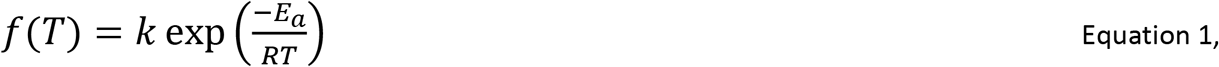

or equivalently,

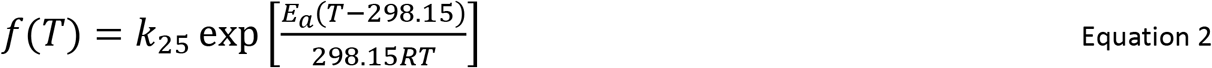

where f(T) is the rate of the process at temperature T, k is a pre-exponential factor, E_a_ is the activation energy in J mol^−1^, R is the universal gas constant of 8.314 J mol^−1^ K^−1^, T in K, k_25_ is the rate of the process at 298.15 K, and 298.15 is the reference temperature in degrees Kelvin (K). As for peaked responses, while a few options are available (Kruse *et al.*, 2008; Hobbs *et al.*, 2013; Heskel *et al.*, 2016), the most commonly implemented version is the modified Arrhenius model of Johnson *et al.* (1942) as presented in Medlyn *et al.* (2002):

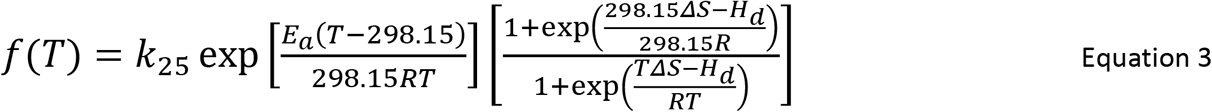

where H_d_ is the deactivation energy in J mol^−1^, and ΔS is the entropy of the process in J mol^−1^ K^−1^.

Equation 3 (**which will be referred to as M2002 from this point forward as it is conventionally referred to in the literature as the source of the equation**) is used for modelling the temperature responses of photosynthetic capacity (i.e. maximum carboxylation capacity of rubisco, V_cmax_, maximum electron transport capacity, J_max_, and triose phosphate utilization capacity (TPU); see Table 1 for a review of which Earth System Models use M2002). These parameters are then used in ecophysiological studies to understand thermal acclimation of photosynthesis (see Kattge & Knorr, 2007; Smith & Dukes, 2017; and Kumarathunge *et al.*, 2019 for examples). Furthermore, this equation is also used in terrestrial biosphere models to predict future carbon cycling (e.g. Rogers *et al.*, 2017).

**Table 1.**
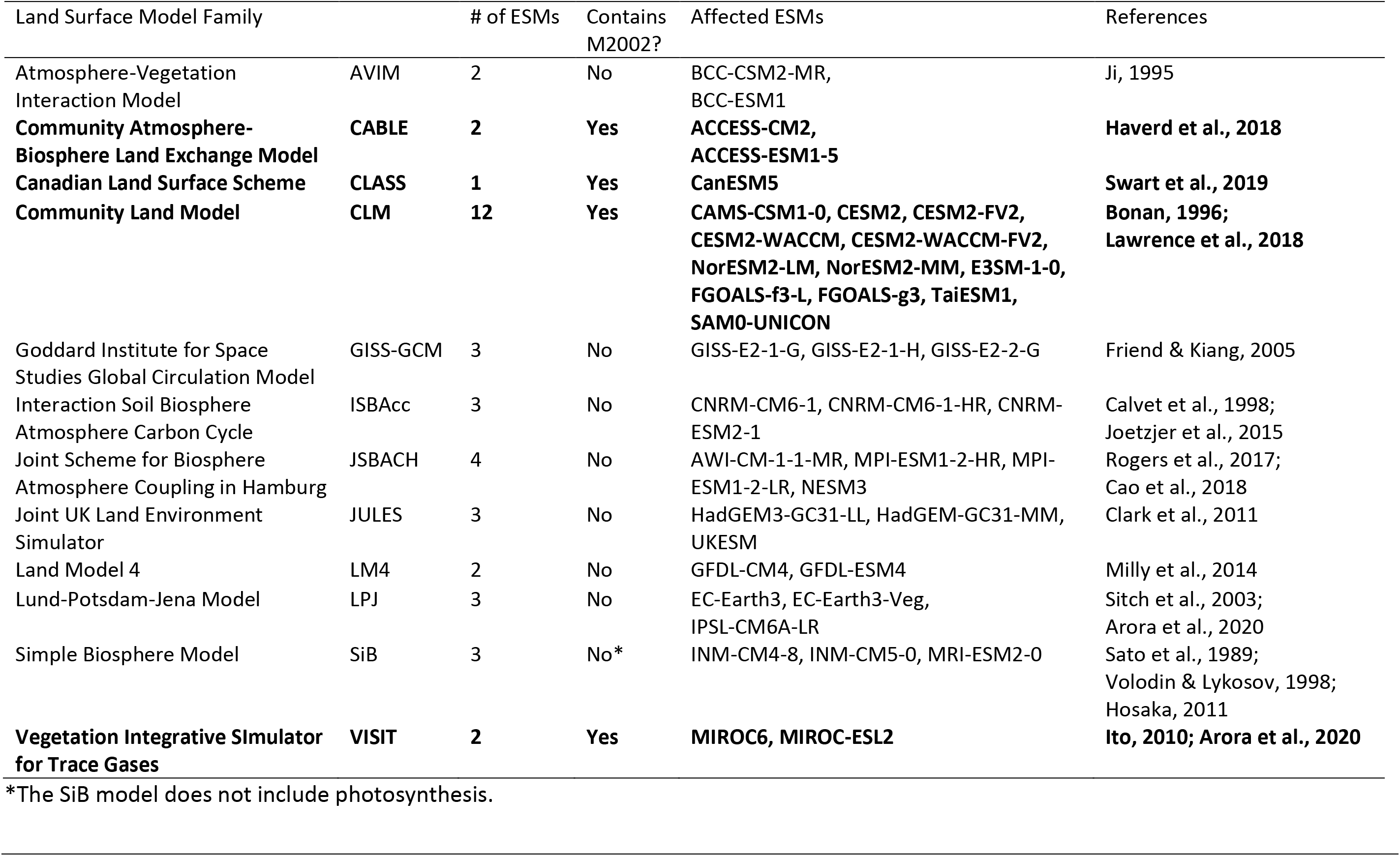
List of land surface models affected by the derivation error in M2002 and the affected Earth Systems Models (ESMs) being used in the CMIP6 simulations. Note that 17 of the 40 (42.5%) CMIP6 ESMs are affected by the derivation error. Affected models are in bold.

Due to the ubiquity of M2002 in modelling temperature responses and the thermal acclimation of photosynthetic capacity (e.g. Kattge & Knorr, 2007; Rogers *et al.*, 2017; Smith & Dukes, 2017; Kumarathunge *et al.*, 2019; Table 1), we revisited the original Johnson *et al.* (1942) modified Arrhenius function to rederive M2002. In the process of this rederivation, we uncovered a term that was completely dropped sometime between Johnson *et al.* (1942) and Medlyn *et al.* (2002) which causes a systematic error in the application of the modified Arrhenius for modelling photosynthetic capacity in individual species (e.g. Medlyn *et al.*, 2002) to modelling global scale carbon uptake (e.g. Rogers *et al.*, 2017; Table 1). We then refit a freely available dataset (Kumarathunge *et al.*, 2019), with both versions of the modified Arrhenius model, and fed the temperature response fits through a carbon balance model to estimate the impact of the derivation error on modelled plant carbon balance. We predicted that the derivation error would cause substantial variation in fitted temperature response parameters, and that these differences would propagate through to modelled daily carbon balance.

## Description

### Rederivation of the modified Arrhenius equation

Johnson *et al.* (1942, equation 24) describe the temperature response of the light intensity of a luciferase reaction whereby the enzyme reversibly denatures or inactivates at high temperatures as:

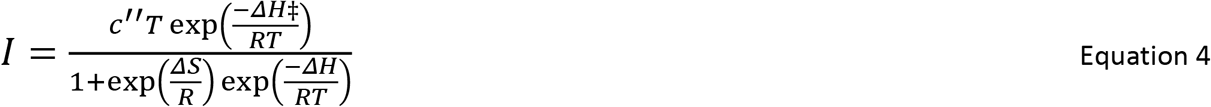

where I is the intensity of the luciferase reaction, c’’ is not explicitly defined in Johnson *et al.* (1942), but appears to represent a constant based on the derivation of Equation 4, T is the temperature in K, R is the universal gas constant of 8.314 J mol^−1^ K^−1^, ΔH‡ is the activation energy in J mol^−1^, ΔH is the deactivation energy in J mol^−1^, and ΔS is the entropy in J mol^−1^ K^−1^. First, we harmonize the notation with standard ecophysiological representations such that ΔH‡ becomes E_a_, and ΔH becomes H_d_:

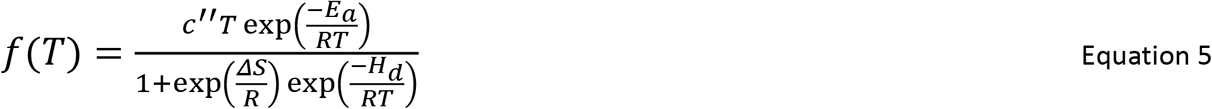

We can relativize the equation to a reference temperature by dividing Equation 5 at a hypothetical temperature by Equation 5 at a standard temperature (i.e. 25 °C):

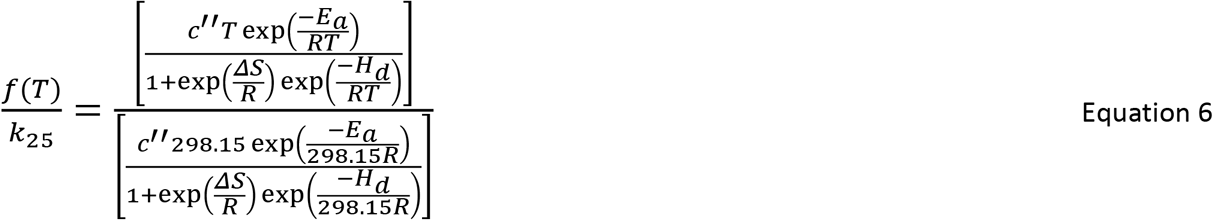

Next, rearrange fractions so that there is one numerator and one denominator on each line for clarity:

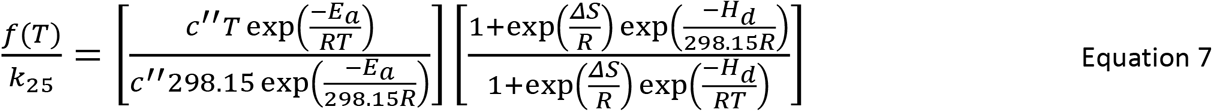

Cancel out the constant c’’, use exponent rules to simplify so that E_a_ is contained within a single exponential function, and the ΔS and H_d_ terms are grouped together:

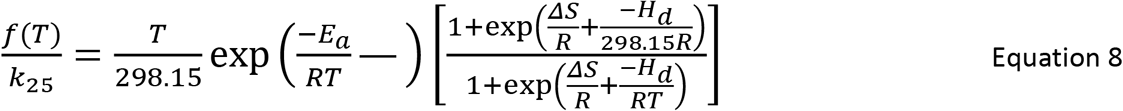

Simplify further by setting a common denominator within exponential terms:

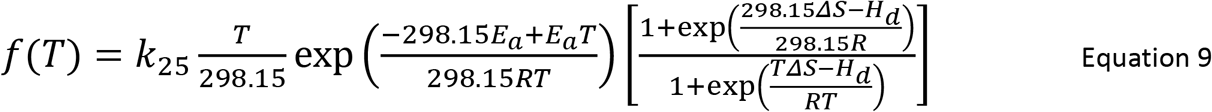

Simplify exponential terms further to reduce the number of instances that E_a_ appears in the equation:

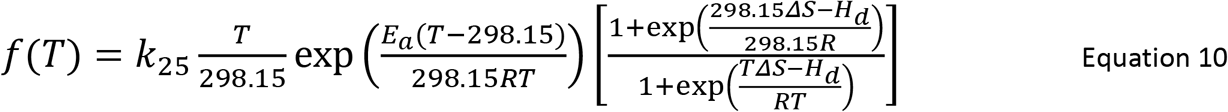

Note the difference between M2002 and the rederived equation 10 (**which will be referred to as J1942**): the term T / 298.15 is missing from M2002. There are several alternative expressions of Equation 4 (e.g. Farquhar *et al*., 1980; Harley *et al.*, 1986; Harley *et al.*, 1992; Harley & Baldocchi, 1995; Lloyd, 1995), however when relativized to a common temperature, the equations presented in Farquhar *et al.* (1980), Harley *et al.* (1992), Harley & Baldocchi (1995), and Lloyd (1995) are all identical to M2002 when relativized by temperature, while the Harley *et al.* (1986) equation is identical to our derivation (J1942) when relativized to a common temperature (see Appendix A for the relativizations of Farquhar *et al.* (1980), Harley *et al.* (1986), and Harley *et al.* (1992); Harley & Baldocchi (1995) uses an identical equation to Eq. 18 from Medlyn *et al.* (2002), while Lloyd (1995) uses an identical equation to Eq. 17 from Medlyn *et al.* (2002)).

We traced the origin of the dropped term to two papers: Hall (1979) and Farquhar et al. (1980). Both papers cite the M2002 equation as belonging to Sharpe and DeMichele (1977) when in fact it belongs to Johnson et al. (1942). In Hall (1979), the reasoning behind the dropped term is given as: “In addition their linear T term was omitted because it has little influence on the function and would have prevented Eq. 22 [the M2002 equation] from approaching Eq. 21 [Equation 1 in the present study] in the limiting conditions of low temperatures” (Hall, 1979, page 305). Considering J1942, at 5 °C the error due to the dropped term would be ~278 K / 298 K = ~0.93, and at 45 °C would be 318 K / 298 K = ~1.07, so ~7% error at those temperatures. In context of measurement precision, such an assumption of “little influence” may have been reasonable at the time. However, the reason regarding the convergence of the two equations is simply tuning the model equations to fit their expectations. Meanwhile, Farquhar et al. (1980) report that: “Eq. (36) is a simplified version of an equation developed by Sharpe and DeMichelle [sic] (1977) to describe the effect of temperature on enzyme inactivation” (Farquhar et al., 1980, page 84). In both cases, it appears that the intention was to simplify the equation, which in the process of simplifying, generated an error that was propagated for over 40 years.

The derivation error introduces multiple systematic errors. First, errors are introduced into the fitted parameters E_a_, H_d_, and ΔS. Second, k_25_ is scaled using the wrong equation, introducing an error in f(T). And third, acclimation equations describing E_a_, H_d_, and ΔS will then be in error due to errors in the fitted parameters at each temperature. Here we focus on the impact of the dropped term on fitted temperature response parameters and modelling whole-plant carbon balance.

### Data analysis

Using A-C_i_ curve data compiled in Kumarathunge *et al.* (2019), available from the Kumarathunge *et al.* (2018) repository, we used the {fitacis} function from the R package {plantecophys} (Duursma, 2015), setting fitmethod = “bilinear”, Tcorrect = FALSE, and fitTPU = TRUE, to obtain V_cmax_ and J_max_. We then fit Equations 3 and 10 to the data, allowing H_d_ to be fit. To ensure that the curves could be fit (i.e. that there were enough data to fit 4 parameters), we only used data where A-C_i_ curves were measured at 5 or more temperatures. This reduced the number of candidate temperature responses from 729 to 403. We used the R package {minpack.lm} (Elzhov *et al.*, 2016), using Equation 2 to obtain starting values for E_a_ and k_25_, while the other initial parameters were ΔS = 0.650 kJ mol^−1^ K^−1^, and H_d_ varying from 1 to 1000 kJ mol^−1^, followed by the {BIC} function to select the best model based on Bayesian Information Criteria. We obtained 341 and 337 successful V_cmax_ temperature response curve fits for M2002 and J1942, respectively, and 241 and 242 successful J_max_ temperature response curve fits for M2002 and J1942, respectively. This resulted in a total of 547 fitted temperature response curves which we filtered further, requiring that E_a_, ΔS, and H_d_ were all positive values and that the V_cmax_ and J_max_ data were paired, resulting in 196 complete pairs of V_cmax_ and J_max_ temperature responses for analysis. The data covered a temperature range from 1 °C to 50 °C across all curves, with a median range of 14 °C to 42.6 °C.

### Modelling

We modelled the impact of the M2002 and J1942 equations on daily net plant carbon balance. Data for leaf area, root and shoot masses, as well as leaf dark respiration at 25 °C were taken for white spruce (*Picea glauca*) from Stinziano & Way (2017), while stomatal conductance model parameters were calculated with the gas exchange data reported in Stinziano & Way (2017). Briefly, Stinziano and Way (2017) grew white spruce in growth chambers under a simulated autumn treatment based on temperatures and photoperiods from Trenton, ON, with weekly harvesting of biomass and gas exchange measurements across 17 weeks as a control treatment, along with a warming treatment where temperatures were +5 °C of the control, a constant summer day/night temperature with declining photoperiod, and a constant summer photoperiod with ambient changes in temperature. Here we only used data from Stinziano & Way (2017) that were from the control treatment. Mean data were taken from the control treatment at weeks 0 and 12 to provide contrasting biomass allocation patterns such that week 0 is a low respiration scenario and week 12 is a high respiration scenario. These different respiration scenarios were used to reduce bias in any conclusions regarding the impact of the M2002 and J1942 equations on carbon balance, as the ratio of photosynthesis to respiration may alter the sensitivity of net carbon balance to the Arrhenius equation used. Root respiration for white spruce was taken from Weger & Guy (1991) and we assumed that stem respiration was equal to root respiration (Table 2).

**Table 2.** Parameters used in modelling daily carbon gain.

| Parameter | Group | Value | Reference |
| --- | --- | --- | --- |
| Respiration | R<sub>dark</sub> | 2.78 μmol m<sup>-2</sup> s<sup>-1</sup> | Stinziano & Way, 2017 |
|  | R<sub>day</sub> | 0.7 * R<sub>dark</sub> | Ayub *et al.*, 2011 |
|  | R<sub>root</sub> | 0.0095 μmol g<sup>-1</sup> s<sup>-1</sup> | Weger & Guy, 1991 |
|  | R<sub>stem</sub> | 0.0095 μmol g<sup>-1</sup> s<sup>-1</sup> | Assumed |
|  | Q<sub>10</sub> | 2.015 | Atkin & Tjoelker, 2003 |
| Γ* | 25 °C | 42.75 μmol mol<sup>-1</sup> | Bernacchi *et al.*, 2001 |
|  | E<sub>a</sub> | 37.83 kJ mol<sup>-1</sup> | Bernacchi *et al.*, 2001 |
| K<sub>m</sub> | 25 °C | 718.4 μmol mol<sup>-1</sup> | Bernacchi *et al.*, 2001 |
|  | E<sub>a</sub> | 65.51 kJ mol<sup>-1</sup> | Bernacchi *et al.*, 2001 |
| α |  | 0.8 | Norman & Campbell, 1998 |
| φ |  | 0.08 | Norman & Campbell, 1998 |
| Leaf Area | Low R | 0.015 m<sup>2</sup> | Stinziano & Way, 2017 |
|  | High R | 0.025 m<sup>2</sup> | Stinziano & Way, 2017 |
| Stem Mass | Low R | 0.496 g | Stinziano & Way, 2017 |
|  | High R | 2.523 g | Stinziano & Way, 2017 |
| Root Mass | Low R | 0.498 g | Stinziano & Way, 2017 |
|  | High R | 5.072 g | Stinziano & Way, 2017 |

**Q<sub>10</sub>: thermal sensitivity coefficient; Γ\*: photorespiratory CO<sub>2</sub> compensation point; K<sub>m</sub>: apparent** **Michaelis-Menten constant for rubisco carboxylation in 21% O<sub>2</sub>/air (i.e. K<sub>c,air</sub>); α: absorbance of** **photosynthetically activation radiation; φ: maximum quantum efficiency of photosynthetic electron** **transport; E<sub>a</sub>: activation energy; R<sub>dark</sub>: leaf respiration in the dark; R<sub>day</sub>: leaf respiration in the light;** **R<sub>root</sub>: root respiration; R<sub>stem</sub>: stem respiration; Low R: low respiration scenario; High R: high** **respiration scenario.**

For the full model structure and equations, please see the accompanying R package {arrhenius.comparison} (“arrhenius.comparison_1.0.2.tar.gz”; Stinziano & Murphy, 2020) (see Table 3 for equations). Briefly, we linked the Medlyn *et al.* (2011) stomatal conductance model with the Farquhar *et al.* (1980) C_3_ photosynthesis model, assuming infinite mesophyll conductance to CO_2_ as these same assumptions were used in fitting the data from Kumarathunge *et al.* (2018). Photosynthetic capacity, both maximum rubisco carboxylation capacity, V_cmax_, and maximum electron transport rate, J_max_, were scaled to temperature using either M2002 or J1942, while respiration was scaled according to (Atkin & Tjoelker, 2003). Photosynthesis and respiration were summed across each modelled day to calculate daily plant carbon assimilation.

**Table 3.**
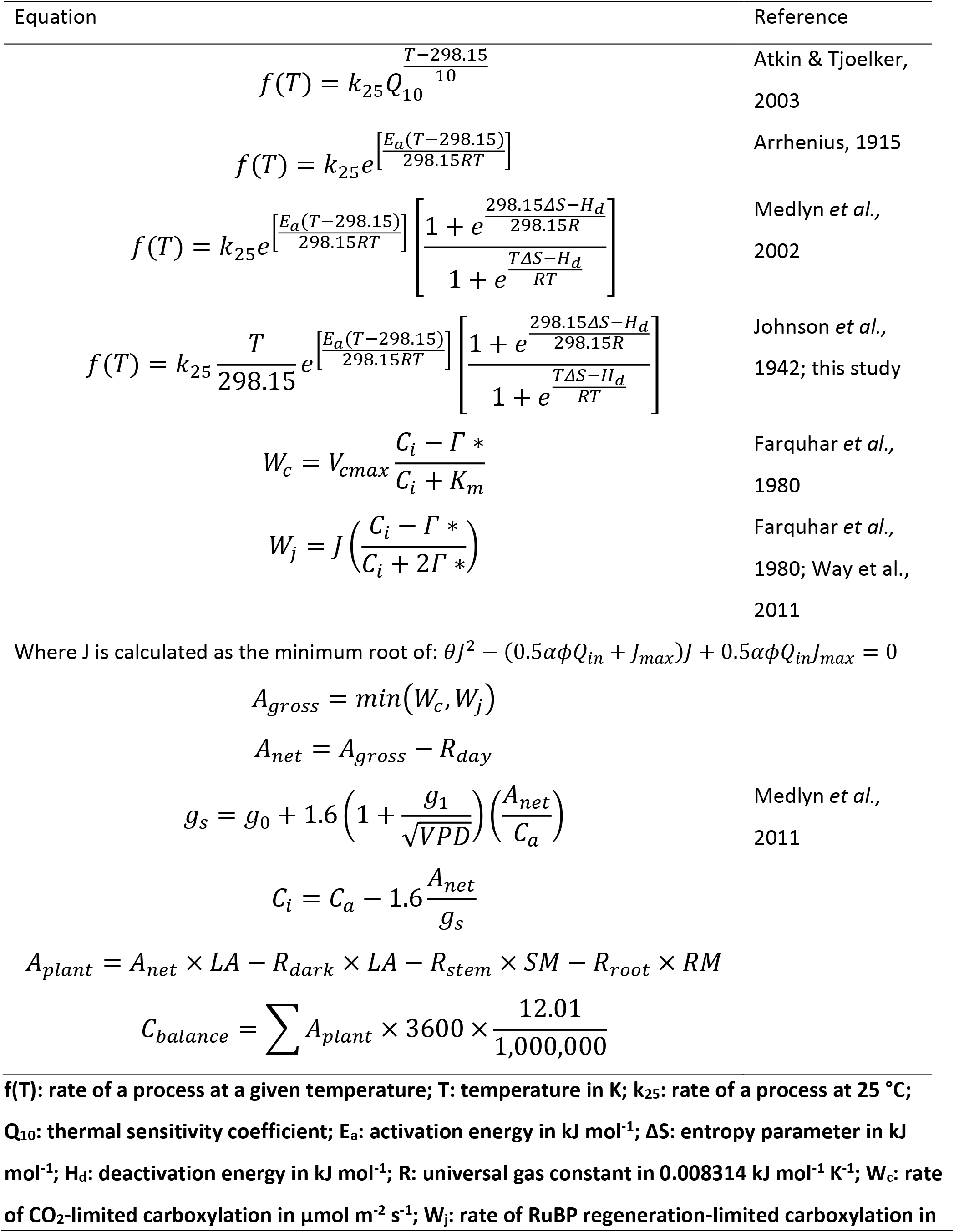
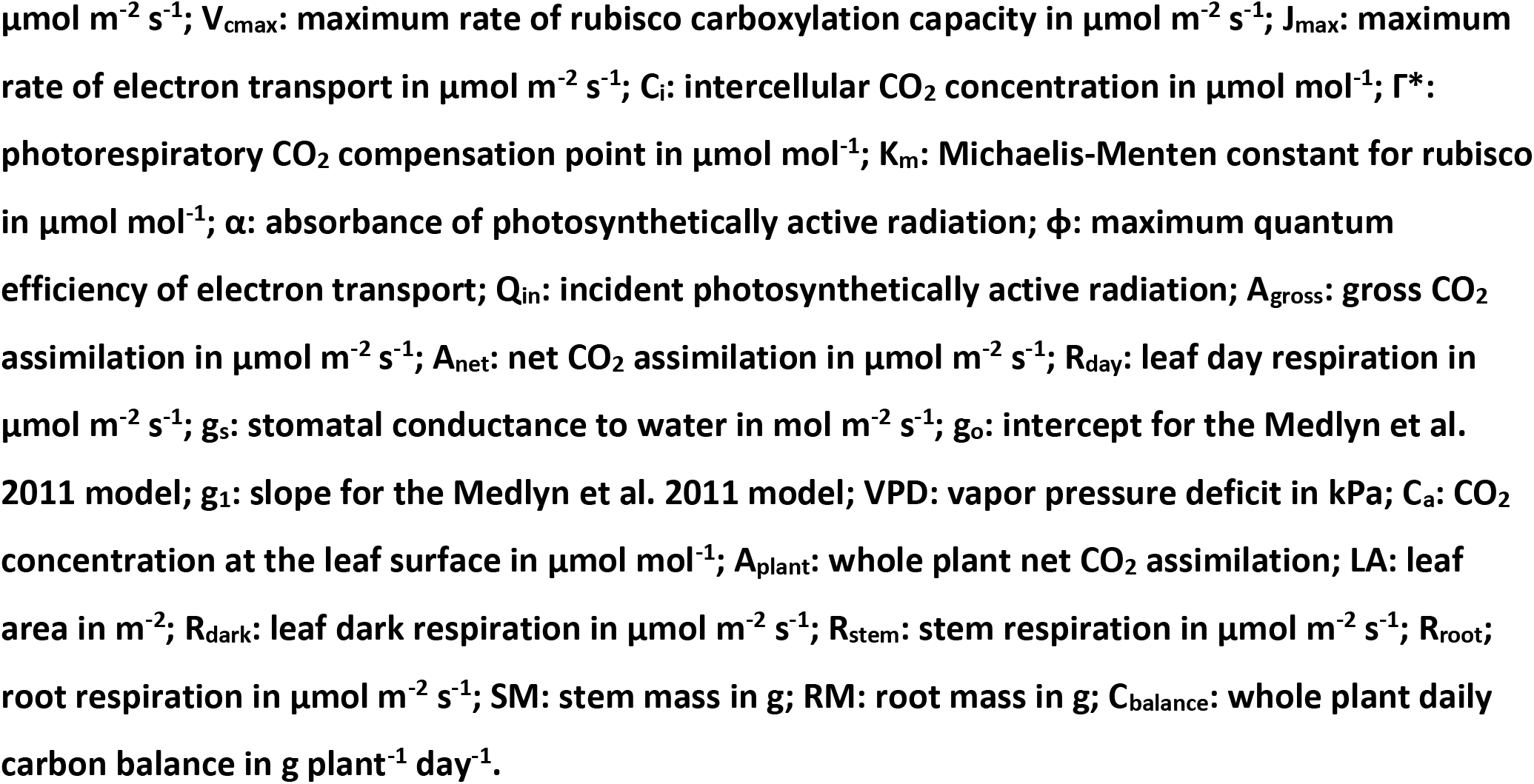
Equations used in modelling daily carbon uptake.

| Equation | Reference |
| --- | --- |
| $f(T) = k_{25} Q_{10}^{\frac{T-298.15}{10}}$ | Atkin & Tjoelker, 2003 |
| $f(T) = k_{25} e^{\left[ \frac{E_a(T-298.15)}{298.15RT} \right]}$ | Arrhenius, 1915 |
| $f(T) = k_{25} e^{\left[ \frac{E_a(T-298.15)}{298.15RT} \right]} \left[ \frac{1 + e^{\frac{298.15\Delta S - H_d}{298.15R}}}{1 + e^{\frac{T\Delta S - H_d}{RT}}} \right]$ | Medlyn *et al.*, 2002 |
| $f(T) = k_{25} \frac{T}{298.15} e^{\left[ \frac{E_a(T-298.15)}{298.15RT} \right]} \left[ \frac{1 + e^{\frac{298.15\Delta S - H_d}{298.15R}}}{1 + e^{\frac{T\Delta S - H_d}{RT}}} \right]$ | Johnson *et al.*, 1942; this study |
| $W_c = V_{cmax} \frac{C_i - \Gamma^*}{C_i + K_m}$ | Farquhar *et al.*, 1980 |
| $W_j = J \left( \frac{C_i - \Gamma^*}{C_i + 2\Gamma^*} \right)$ | Farquhar *et al.*, 1980; Way *et al.*, 2011 |
| Where J is calculated as the minimum root of: $\theta J^2 - (0.5\alpha\phi Q_{in} + J_{max})J + 0.5\alpha\phi Q_{in}J_{max} = 0$ | |
| $A_{gross} = \min(W_c, W_j)$ | |
| $A_{net} = A_{gross} - R_{day}$ | |
| $g_s = g_0 + 1.6 \left( 1 + \frac{g_1}{\sqrt{VPD}} \right) \left( \frac{A_{net}}{C_a} \right)$ | Medlyn *et al.*, 2011 |
| $C_i = C_a - 1.6 \frac{A_{net}}{g_s}$ | |
| $A_{plant} = A_{net} \times LA - R_{dark} \times LA - R_{stem} \times SM - R_{root} \times RM$ | |
| $C_{balance} = \sum A_{plant} \times 3600 \times \frac{12.01}{1,000,000}$ | |
| 346 **f(T): rate of a process at a given temperature; T: temperature in K; k<sub>25</sub>: rate of a process at 25 °C;**<br>347 **Q<sub>10</sub>: thermal sensitivity coefficient; E<sub>a</sub>: activation energy in kJ mol<sup>-1</sup>; ΔS: entropy parameter in kJ**<br>348 **mol<sup>-1</sup>; H<sub>d</sub>: deactivation energy in kJ mol<sup>-1</sup>; R: universal gas constant in 0.008314 kJ mol<sup>-1</sup> K<sup>-1</sup>; W<sub>c</sub>: rate**<br>349 **of CO<sub>2</sub>-limited carboxylation in μmol m<sup>-2</sup> s<sup>-1</sup>; W<sub>j</sub>: rate of RuBP regeneration-limited carboxylation in** |  |

Modelling was performed on 18 total days of environmental data, with three days of data from three months (17^th^ – 19^th^ of May, August, and October, 2019) obtained from external irradiance sensors at the Biotron Experimental Climate Change Research Centre at the University of Western Ontario and the remaining environmental data from Environment Canada historical climate data for South London (43.01°N, 81.27°W, altitude: 251 m; temperature range: −0.1 – 27.9 °C) and the rooftop greenhouse at the University of New Mexico (35.08°N, 106.62°W, altitude: 1587 m; temperature range: 7.2 – 42.2 °C) to capture different levels of environmental variability (see *Impacts on modelled net carbon balance* for environmental data).

Overall, the modelling approach allows us to assess the relative differences between Equations 3 and 10 under a low- and high- respiration scenario across different ranges of seasonal variability.

#### Statistical analysis

Data were analyzed using the {lm} function in R v.3.6.2 (R Core Team, 2019), regressing the data obtained from M2002 against the data obtained from J1942 for each of V_cmax,25_, J_max,25_, E_a,Vcmax_, E_a,Jmax_, ΔS_Vcmax_, ΔS_Jmax_, H_d,Vcmax_, H_d,Jmax_, BIC values for the fits of Equations 3 and 10, daily photosynthesis, daily net carbon balance, and daily photosynthesis : daily respiration ratios. Intercepts in the regression, when significant, were interpreted as a bias in the parameter, while deviations in slope from a 1:1 relationship where interpreted as percentage over- or under-estimation of the parameter (i.e. [parameter slope – 1] ∙ 100 = % estimation error). All p-values were corrected for multiple testing using the p.adjust() function in R with Holm’s method. All code and data will be made freely available on GitHub upon publication in the {arrhenius.comparison} R package (Stinziano & Murphy, 2020).

## Results

### Equations M2002 and J1942 exhibited similar performance

The performance between M2002 and J1942 for fitting V_cmax,25_ temperature responses were essentially identical when assessed based on BIC, with a slope of 1.001 ± 0.001 (*F*_*1,195*_ = 5.52 ∙ 10^5^, *R*^*2*^ = 0.9996, *P* < 2.2 ∙ 10^−16^; Fig. **1a**), as was the case for J_max,25_ with a slope of 1.001 ± 0.001 (*F*_*1,195*_ = 2.43 ∙ 10^6^, *R*^*2*^ = 0.9999, *P* < 2.2 ∙ 10^−16^; Fig. **1b**). However, while fitted temperature responses based on BIC appeared identical, the largest discrepancy between fitted responses was observed when comparing J_max,25_ of both plant species scaled across 10 °C and 50 °C (Figs. **1c-f**; note the differences between the blue and red hatched lines).

**Figure 1.**
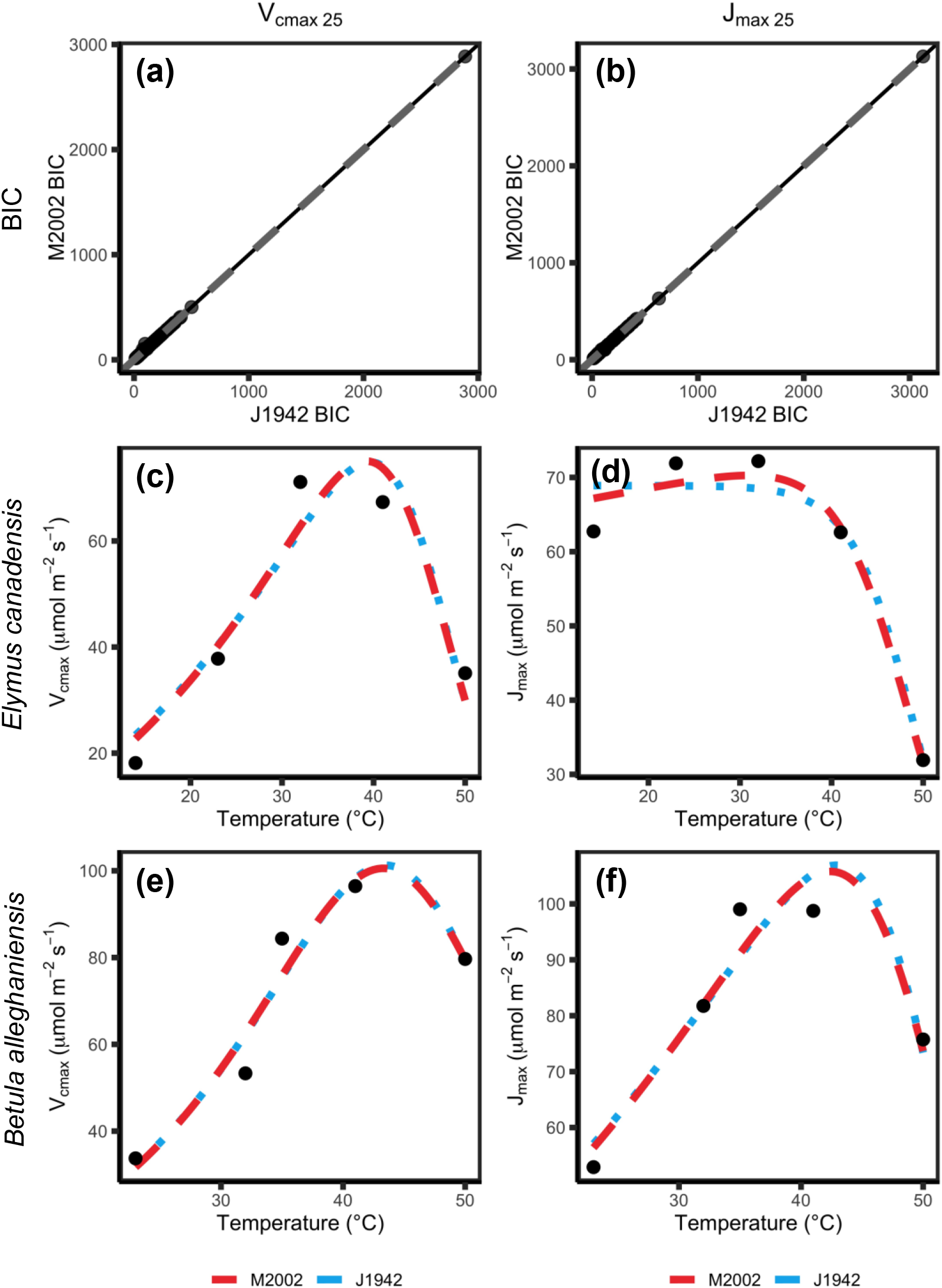
Relative performance and fit for Equations 3 (M2002) and 10 (J1942) for temperature responses of V_cmax_ 25 and J_max 25_ based on a,b) Bayesian Information Criterion (BIC), and c-f) visual inspection of curve fits across 10-50 °C for c,d) *Elymus canadensis* and 20-50 °C of e,f) *Betula alleghaniensis*. Red dashed lines indicate curves fitted with the M2002 equation and blue dotted lines indicate curves fitted with the J1942 equation. Parameters for c-f) are as follows: c) M2002: k_25_: 46.4 μmol m^−2^ s^−1^, E_a_: 29.0 kJ mol^−1^, ΔS: 0.692 kJ mol^−1^ K^−1^, H_d_: 221.4 kJ mol^−1^; J1942: k_25_: 46.4 μmol m^−2^ s^−1^, E_a_: 29.0 kJ mol^−1^, ΔS: 0.655 kJ mol^−1^ K^−1^, H_d_: 209.0 kJ mol^−1^; d) M2002: k_25_: 68.9 μmol m^−2^ s^−1^, E_a_: 0.0 kJ mol^−1^, ΔS: 0.746 kJ mol^−1^ K^−1^, H_d_: 240.5 kJ mol^−1^; J1942: k_25_: 69.5 μmol m^−2^ s^−1^, E_a_: 0.0 kJ mol^−1^, ΔS: 0.625 kJ mol^−1^ K^−1^, H_d_: 201.0 kJ mol^−1^; e) M2002: k_25_: 37.6 μmol m^−2^ s^−1^, E_a_: 57.7 kJ mol^−1^, ΔS: 0.579 kJ mol^−1^ K^−1^, H_d_: 185.4 kJ mol^−1^; J1942: k_25_: 37.0 μmol m^−2^ s^−1^, E_a_: 57.7 kJ mol^−1^, ΔS: 0.557 kJ mol^−1^ K^−1^, H_d_: 178.1 kJ mol^−1^; f) M2002: k_25_: 66.7 μmol m^−2^ s^−1^, E_a_: 23.0 kJ mol^−1^, ΔS: 0.761 kJ mol^−1^ K^−1^, H_d_: 246.6 kJ mol^−1^; J1942: k_25_: 65.5 μmol m^−2^ s^−1^, E_a_: 23.0 kJ mol^−1^, ΔS: 0.668 kJ mol^−1^ K^−1^, H_d_: 216.1 kJ mol^−1^.

Estimates of V_cmax,25_ were essentially identical between the M2002 fitting (y-axis) and the J1942 fitting (x-axis) with a slope of 1.002 ± 0.004 (*F*_*1,195*_ = 5.84 ∙ 10^4^, *R*^*2*^ = 0.9967, *P* < 2.2 ∙ 10^−16^; Fig. **2e**), as was the case for J_max,25_ with a slope of 1.001 ± 0.001 (*F*_*1,195*_ = 6.56 ∙ 10^5^, *R*^*2*^ = 0.9997, *P* < 2.2 ∙ 10^−16^; Fig. **2f**). E_a,Vcmax_ was generally underestimated with a slope of 0.847 ± 0.024 and a positive bias of 9.73 ± 2.59 kJ mol^−1^ when using M2002 (*F*_*1,194*_ = 1202, *R*^*2*^ = 0.8610, *P* < 2.2 ∙ 10^−16^; Fig. **2a**), while E_a,Jmax_ was underestimated with a slope of 0.832 ± 0.013 and a positive bias of 5.98 ± 1.06 kJ mol^−1^ (*F*_*1,194*_ = 3834, *R*^*2*^ = 0.9518, *P* < 2.2 ∙ 10^−16^; Fig. **2b**). ΔS_Vcmax_ was essentially identical between M2002 and J1942 with a slope of 1.000 ± 0.0233 (*F*_*1,195*_ = 1849, *R*^*2*^ = 0.9046, *P* < 2.2 ∙ 10^−16^; Fig. **2c**), as was the case with ΔS_Jmax_ with a slope of 0.988 ± 0.033 (*F*_*1,195*_ = 904, *R*^*2*^ = 0.8226, *P* < 2.2 ∙ 10^−16^; Fig. **2d**), and H_d,Vcmax_ with a slope of 1.002 ± 0.005 (*F*_*1,195*_ = 4.12 ∙ 10^4^, *R*^*2*^ = 0.9953, *P* < 2.2 ∙ 10^−16^; Fig. **2g**). However, H_d,Jmax_ was underestimated with a slope of 0.952 ± 0.024 and a positive bias of 31.48 ± 8.93 kJ mol^−1^ (*F*_*1,194*_ = 1592, *R*^*2*^ = 0.8914, *P* < 2.2 ∙ 10^−16^; Fig. **2h**).

**Figure 2.**
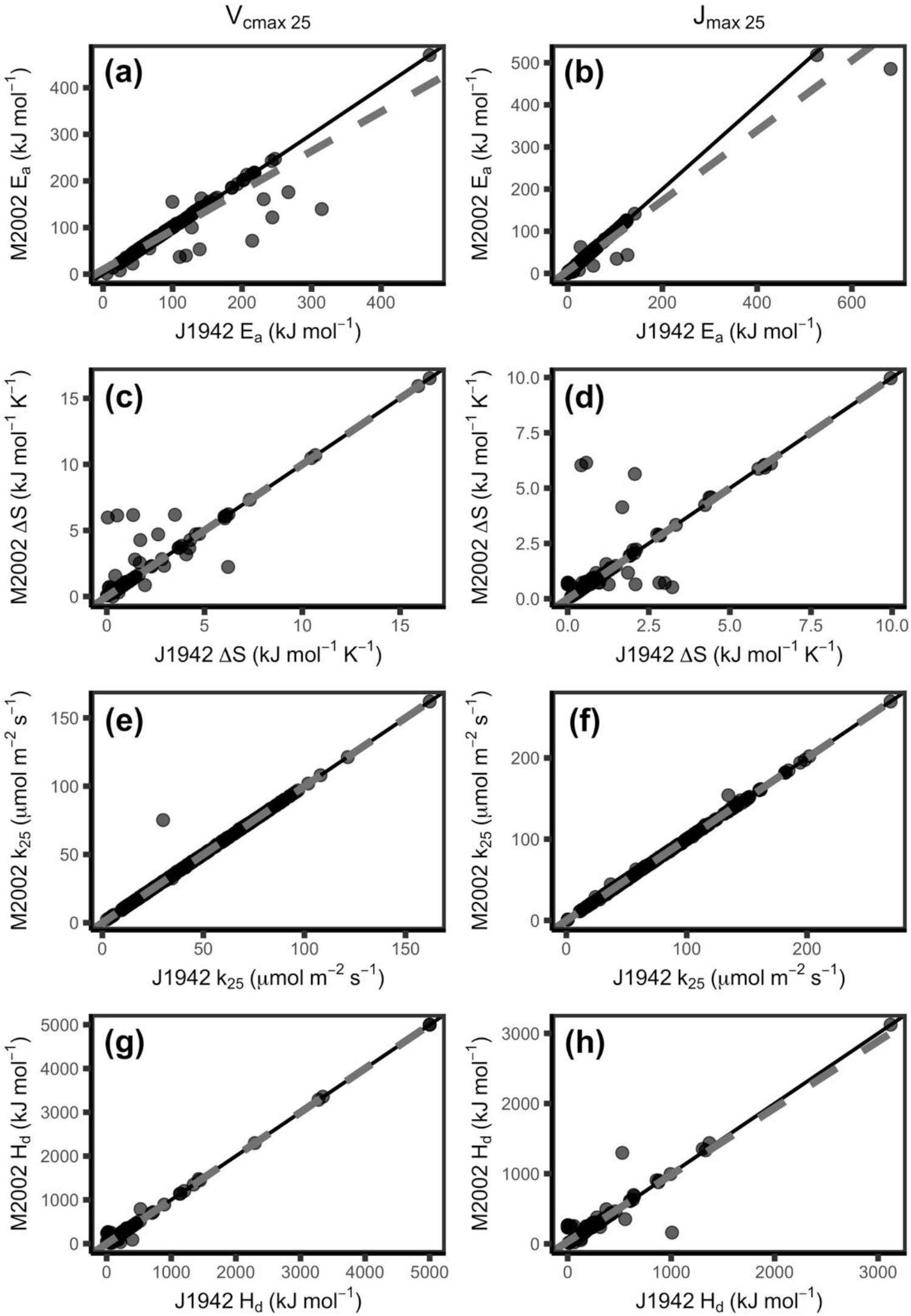
The modified Arrhenius equation missing the term (M2002) compared to J1942 fits different values for E_a_ (a,b), but mostly similar values for ΔS (c,d), k_25_ (e,f) and H_d_ (f, g) for both V_cmax_ (a, c, e, g) and J_max_ (c, d, f, h). E_a_: activation energy, ΔS: entropy parameter, k_25_: rate of the process at 25 °C, H_d_: deactivation energy, V_cmax_: maximum capacity of rubisco carboxylation, J_max_: maximum rate of electron transport. Black line indicates 1:1 line and grey dashed line indicates respective modelled slopes and intercepts.

### Impacts on modelled net carbon balance

In general, the differences in thermal response parameters were amplified when integrated at the whole-plant level across a range of environmental conditions (Fig. **3**). For modelled daily photosynthesis (A), the slope for the low respiration model was 0.819 ± 0.013 and the intercept was 0.018 ± 0.002 g plant^−1^ day^−1^ (M2002 versus J1942; approximately 18% of modelled A) (*F*_1,3523_ = 4.168 ∙ 10^3^, R^2^ = 0.5418; *P* < 2.2 ∙ 10^−16^) (Fig. **4a**). For the high respiration model of daily A, the slope was 0.819 ± 0.013 and the intercept was 0.031 ± 0.003 g plant^−1^ day^−1^ (approximately 18% of modelled A) (*F*_1,3523_ = 4.168 ∙ 10^3^, R^2^ = 0.5418; *P* < 2.2 ∙ 10^−16^) (Fig. **4b**). The low respiration model of total daily carbon (C) gain had a slope of 0.836 ± 0.012 and an intercept of 0.012 ± 0.001 g plant^−1^ day^−1^ (approximately 16% of modelled C gain) (M2002 versus J1942; *F*_1,3523_ = 4.539 ∙ 10^3^, R^2^ = 0.5629; *P* <2.2 ∙ 10^−16^) (Fig. **4c**). The high respiration of total C gain similarly had a slope of 0.859 ± 0.012 and an intercept of 0.011 ± 0.002 g plant^−1^ day^−1^ (approximately 14% of modelled C gain) (M2002 versus J1942; *F*_1,3523_ = 5.364 ∙ 10^3^, R^2^ = 0.6035; *P* <2.2 ∙ 10^−16^) (Fig. **4d**). The ratio of the total daily photosynthesis: respiration (A/R) was also considered when comparing models. The low respiration model of A/R had a slope of 0.982 ± 0.007 and the intercept was 0.146 ± 0.031 (approximately 2% of modelled A/R) (M2002 versus J1942; *F*_1,3523_ = 2.096 ∙ 10^4^, R^2^ = 0.8561; *P* <2.2 ∙ 10^−16^) (Fig. **4e**). The high respiration model had a similar slope of 0.982 ± 0.007 and the intercept was 0.082 ± 0.017 (approximately 2% of modelled A/R) (M2002 versus J1942; *F*_1,3523_ = 2.097 ∙ 10^4^, R^2^ = 0.8562; *P* <2.2 ∙ 10^−16^) (Fig. **4f**).

**Figure 3.**
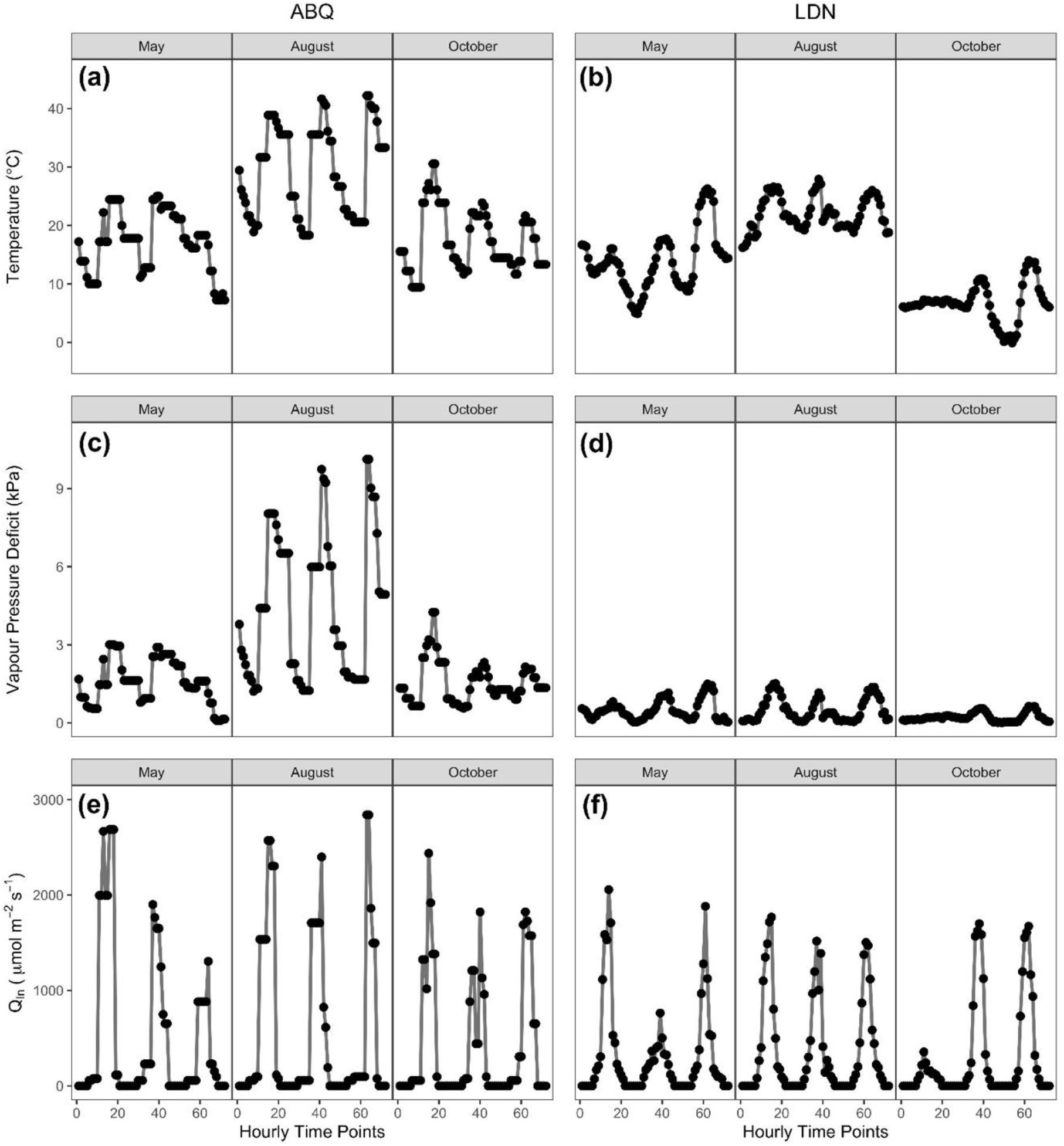
Hourly environmental data used to drive the model in Table 3 covering 3 days (17^th^, 18^th^, and 19^th^) of 3 months. (a,c,e) Albuquerque, NM, USA (ABQ); (b,d,e) London, ON, Canada (LDN). Environmental parameters include: a,b) temperature; c,d) vapour pressure deficit; and e,f) irradiance (Q_in_).

**Figure 4.**
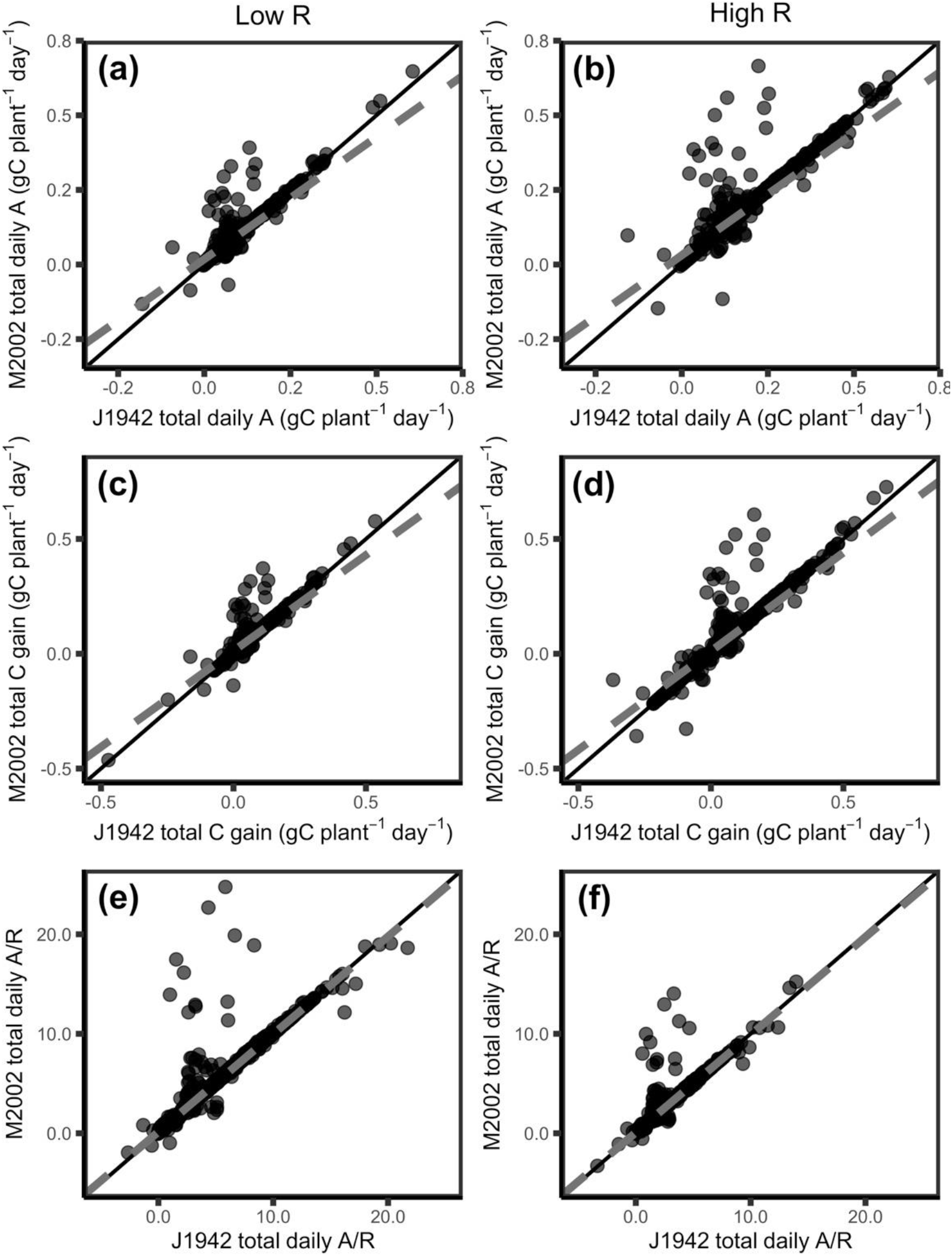
The modified Arrhenius equation without the missing term (M2002) gives differences in modelled daily carbon fluxes compared to J1942 for low R (a,c,e) and high R (b,d,f) under scenarios where H_d_ is allowed to vary. Modelled carbon fluxes include: a,b) Total daily photosynthesis; c,d) total daily C gain; and e,f) total daily A/R. A: photosynthesis, R respiration, Low R: low respiration, High R: high respiration. Black line indicates 1:1 line and grey dashed line indicates respective modelled slopes and intercepts.

## Discussion

We sought to determine whether the missing term in M2002 had a meaningful impact on fitted temperature response parameters due to its prevalence in photosynthetic temperature response data and vegetation modelling (Kattge & Knorr, 2007; Duursma & Medlyn, 2012; Rogers *et al.*, 2017; Smith & Dukes, 2017; Stinziano *et al.*, 2018; Stinziano *et al.*, 2019; Kumarathunge *et al.*, 2019). Our present analysis suggests that there is a large impact on E_a_ for both V_cmax_ and J_max_, however there were minimal impacts on k_25_, H_d_, and ΔS (though note that H_d,Jmax_ was underestimated) (Fig. **2**). In general, fitting M2002 instead of J1942 results in E_a_ reductions of around 15% with positive bias, even though fitted responses are *visually* similar (Fig. **1**). These findings are promising in that one of the parameters to which modelled carbon gain is particularly sensitive, H_d_ (Stinziano *et al.*, 2018), is minimally affected by the missing term. However, since temperature responses are non-linear, small changes in the shape of an accelerating curve can have a strong impact on the integral of the response (Jensen, 1906; e.g. changes in the shape of the temperature response of carbon assimilation can have strong impacts on total carbon fixation). Despite fit performance being nearly identical and visually similar (Fig. **1**), the differences due to E_a_ values between the equations led to an 18% reduction in modelled daily photosynthesis for M2002 compared to J1942, leading to a ~14-16% reduction in modelled net daily carbon gain. Overall, comparisons of low-respiration to high-respiration scenarios resulted in similar percent changes in daily C balance with similar variation.

Not only did we compare the impact of the missing term in M2002 on different respiration scenarios, but we also modelled data across a range of environmental conditions to understand the impact on C balances of plants under different temperature regimes and how these impacts could scale up in global models. A ~15% reduction in net daily carbon gain is substantial by itself, but this difference would also be amplified over time for a single plant. Plant growth follows the compound interest law (Blackman, 1919). As leaf area increases with growth so too does the rate of photosynthesis and thus the amount of carbon available to further increase growth and metabolism, hence why relative growth rates are commonly utilized in the literature (e.g. Shipley, 1989; Causton, 1991; Tjoelker *et al.*, 1999; Loveys *et al.*, 2003; Poorter *et al.*, 2012; Pommereng & Anders, 2015). A reduction of 18% in modelled daily photosynthesis caused by using the modified equation with the missing term (M2002) would become compounded long-term with plant growth and may result in underestimations of future carbon uptake. It is thus likely that the differences we observed would accumulate to even larger carbon flux errors across large spatial and temporal scales with fluctuating temperatures.

Based on the above analysis, the impact of the missing term in the modified Arrhenius equation substantially alters E_a_ and net daily carbon balance at a whole-plant level. While our models of plant carbon uptake were indicative of problems caused by using M2002, it would be worth comparing how the use of this derivation has affected the interpretation of temperature response data and whole plant carbon balances collected from previous studies. Future work comparing the impact of using M2002 opposed to J1942 on interpreting whole tree carbon balance and net primary production data could lead to better estimations of the temperature sensitivity of carbon uptake. Given that carbon balance is the time integral of net CO_2_ assimilation, this may also lead to substantial impacts over a long time period. On a larger scale, a correction of this derivation error will also be necessary when considering Earth system models, as 17 out of 40 models in the CMIP6 simulations are affected by the derivation error (Table 1). We therefore recommend the switch from M2002 to J1942 because: 1) M2002 is the result of a derivation error circa 1979, and 2) M2002 and J1942 lead to different net daily carbon estimates due to the derivation error, which may currently be compensated by other factors in large-scale models, most notably in the Community Land Model which currently relies on the M2002 equation (Bonan, 1996; Lawrence et al., 2018) and underlies 30% of Earth System Models (Table 1).

## Acknowledgments

We would like to thank Wesley J. Noe at the University of New Mexico for providing climate data. This research was supported by personal funds.

## Author Contributions

Both authors contributed to all aspects of the study.

## Additional Information

A version of this manuscript was posted on bioRxiv (manuscript ID: BIORXIV/2020/921973).

## Supplementary information

## Appendix A Alternate derivations of the modified Arrhenius equation

Below are the temperature relativizations of Farquhar et al. (1980), Harley et al. (1986), and Harley et al. (1992) showing that Farquhar et al. (1980) and Harley et al. (1992) can be transformed to be identical to Equation 3 in the manuscript, while Harley et al. (1986) can be transformed to be identical to Equation 10 in the manuscript.

*Farquhar et al. (1980)*

From Eq. 36 in Farquhar et al. (1980):

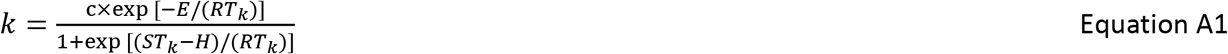

Relativizing to 25 °C:

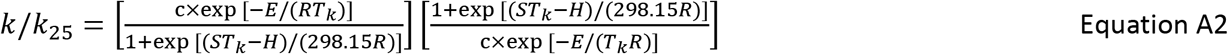

Simplifying:

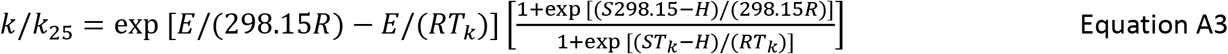

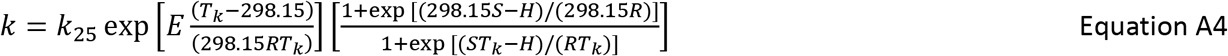

where E is activation energy (E_a_), S is deactivation entropy (ΔS), and H is deactivation energy (H_d_).

*Harley et al. (1986)*

From Eq. 7 in Harley et al. (1986):

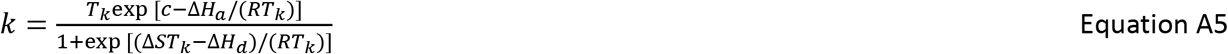

Relativizing to 25 °C:

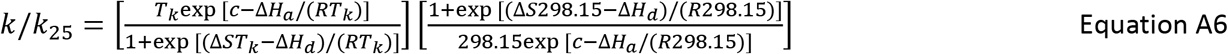

Simplifying:

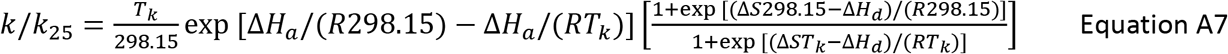

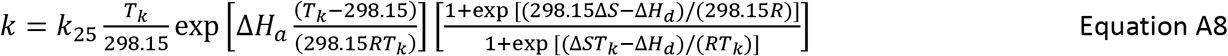

where ΔH_a_ is activation energy (E_a_), and ΔH_d_ is deactivation energy (H_d_).

*Harley et al. (1992)*

From Eq. 9 in Harley et al. (1992):

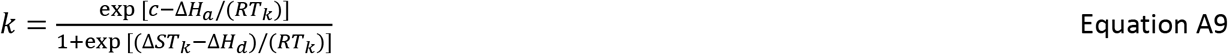

Relativizing to 25 °C:

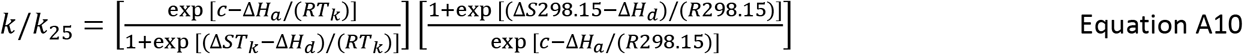

Simplifying:

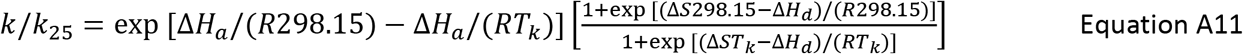

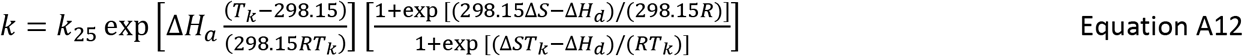

where ΔH_a_ is activation energy (E_a_), and ΔH_d_ is deactivation energy (H_d_).

