## Supplementary Information/Appendix for "A derivation error that affects carbon balance models exists in the current implementation of the modified Arrhenius function" (from another version of this preprint: version 2, 2020-04-23)

**Running Title: Derivation error in modified Arrhenius model**

Author: Bridget Murphy[1], Joseph R. Stinziano[2,*]

Affiliations: [1] Department of Biology, University of Western Ontario, London, ON, Canada;
[2] Department of Biology, University of New Mexico, Albuquerque, NM, USA

### Supplementary information

### Appendix A – Alternate derivations of the modified Arrhenius equation

Below are the temperature relativizations of Farquhar et al. (1980), Harley et al. (1986), and Harley et al. (1992) showing that Farquhar et al. (1980) and Harley et al. (1992) can be transformed to be identical to Equation 3 in the manuscript, while Harley et al. (1986) can be transformed to be identical to Equation 10 in the manuscript.

*Farquhar et al. (1980)*

From Eq. 36 in Farquhar et al. (1980):

$$k = \frac{\mathrm{c}\times\exp\left[-E/(RT_k)\right]}{1+\exp\left[(ST_k-H)/(RT_k)\right]} \qquad \text{Equation A1}$$

Relativizing to 25 °C:

$$k/k_{25} = \left[\frac{\mathrm{c}\times\exp\left[-E/(RT_k)\right]}{1+\exp\left[(ST_k-H)/(298.15R)\right]}\right]\left[\frac{1+\exp\left[(ST_k-H)/(298.15R)\right]}{\mathrm{c}\times\exp\left[-E/(T_kR)\right]}\right] \qquad \text{Equation A2}$$

Simplifying:

$$k/k_{25} = \exp\left[E/(298.15R) - E/(RT_k)\right]\left[\frac{1+\exp\left[(S298.15-H)/(298.15R)\right]}{1+\exp\left[(ST_k-H)/(RT_k)\right]}\right] \qquad \text{Equation A3}$$

$$k = k_{25}\exp\left[E\frac{(T_k-298.15)}{(298.15RT_k)}\right]\left[\frac{1+\exp\left[(298.15S-H)/(298.15R)\right]}{1+\exp\left[(ST_k-H)/(RT_k)\right]}\right] \qquad \text{Equation A4}$$

where E is activation energy ($E_a$), S is deactivation entropy ($\Delta S$), and H is deactivation energy ($H_d$).

*Harley et al. (1986)*

From Eq. 7 in Harley et al. (1986):

$$k = \frac{T_k \exp\left[c - \Delta H_a/(RT_k)\right]}{1 + \exp\left[(\Delta S T_k - \Delta H_d)/(RT_k)\right]} \qquad \text{Equation A5}$$

Relativizing to 25 °C:

$$k/k_{25} = \left[\frac{T_k \exp\left[c - \Delta H_a/(RT_k)\right]}{1 + \exp\left[(\Delta S T_k - \Delta H_d)/(RT_k)\right]}\right]\left[\frac{1 + \exp\left[(\Delta S 298.15 - \Delta H_d)/(R298.15)\right]}{298.15 \exp\left[c - \Delta H_a/(R298.15)\right]}\right] \qquad \text{Equation A6}$$

Simplifying:

$$k/k_{25} = \frac{T_k}{298.15} \exp\left[\Delta H_a/(R298.15) - \Delta H_a/(RT_k)\right]\left[\frac{1 + \exp\left[(\Delta S 298.15 - \Delta H_d)/(R298.15)\right]}{1 + \exp\left[(\Delta S T_k - \Delta H_d)/(RT_k)\right]}\right] \qquad \text{Equation A7}$$

$$k = k_{25} \frac{T_k}{298.15} \exp\left[\Delta H_a \frac{(T_k - 298.15)}{(298.15 R T_k)}\right]\left[\frac{1 + \exp\left[(298.15 \Delta S - \Delta H_d)/(298.15 R)\right]}{1 + \exp\left[(\Delta S T_k - \Delta H_d)/(RT_k)\right]}\right] \qquad \text{Equation A8}$$

where $\Delta H_a$ is activation energy ($E_a$), and $\Delta H_d$ is deactivation energy ($H_d$).

*Harley et al. (1992)*

From Eq. 9 in Harley et al. (1992):

$$k = \frac{\exp\left[c - \Delta H_a/(RT_k)\right]}{1 + \exp\left[(\Delta S T_k - \Delta H_d)/(RT_k)\right]} \qquad \text{Equation A9}$$

Relativizing to 25 °C:

$$k/k_{25} = \left[\frac{\exp\left[c - \Delta H_a/(RT_k)\right]}{1 + \exp\left[(\Delta S T_k - \Delta H_d)/(RT_k)\right]}\right]\left[\frac{1 + \exp\left[(\Delta S 298.15 - \Delta H_d)/(R298.15)\right]}{\exp\left[c - \Delta H_a/(R298.15)\right]}\right] \qquad \text{Equation A10}$$

Simplifying:

$$k/k_{25} = \exp\left[\Delta H_a/(R298.15) - \Delta H_a/(RT_k)\right]\left[\frac{1 + \exp\left[(\Delta S 298.15 - \Delta H_d)/(R298.15)\right]}{1 + \exp\left[(\Delta S T_k - \Delta H_d)/(RT_k)\right]}\right] \qquad \text{Equation A11}$$

$$k = k_{25} \exp\left[\Delta H_a \frac{(T_k - 298.15)}{(298.15 R T_k)}\right]\left[\frac{1 + \exp\left[(298.15 \Delta S - \Delta H_d)/(298.15 R)\right]}{1 + \exp\left[(\Delta S T_k - \Delta H_d)/(RT_k)\right]}\right] \qquad \text{Equation A12}$$

where $\Delta H_a$ is activation energy ($E_a$), and $\Delta H_d$ is deactivation energy ($H_d$).

**Supplementary Information**

*Instructions for installing {arrhenius.comparison} in R for review purposes*

1. Download the arrhenius.comparison.tar.gz file
2. Set working directory in R to the directory that holds the arrhenius.comparison.tar.gz file using setwd()
3. Run:

```
install.packages("arrhenius.comparison_1.0.1.tar.gz", repos = NULL, type = "source")
library(arrhenius.comparison)
```
