## Supplementary material for "A derivation error that affects carbon balance models exists in the current implementation of the modified Arrhenius function": R package for analysis: Modeling_carbon_balance_BM.html (from another version of this preprint: version 2, 2020-04-23)


### Modeling\_carbon\_balance

###### Joseph R. Stinziano

###### `r Sys.Date()

```
library(arrhenius.comparison)
library(dplyr)
```

```
## 
## Attaching package: 'dplyr'
```

```
## The following objects are masked from 'package:stats':
## 
##     filter, lag
```

```
## The following objects are masked from 'package:base':
## 
##     intersect, setdiff, setequal, union
```

Basically need to run week 1 and week 2 code for environmental data and compile Note that the output will include a sum across all the environmental data in the input file. We will need to add descriptors for each day and then sum by day. As long as DOY is specified in the environment data, we will be able to do this easily since calculate\_photo binds the environmental data to the physiology data.

#Hd = fitted #Week 1 modeling - low root/leaf ratio #Remove octothorpe below to run chunk #```{r include = FALSE} data\_env <- read.csv(system.file(“extdata”, “env\_data.csv”, package = “arrhenius.comparison”), stringsAsFactors = FALSE) pars\_v <- read.csv(system.file(“extdata”, “fits\_v\_3.csv”, package = “arrhenius.comparison”), stringsAsFactors = FALSE) pars\_j <- read.csv(system.file(“extdata”, “fits\_j\_3.csv”, package = “arrhenius.comparison”), stringsAsFactors = FALSE) pars\_v <- pars\_v[pars\_v$Hd != 200,] pars\_j <- pars\_v[pars\_j$Hd != 200,] data\_phys <- list(NULL) #Basically, we need a solid set of respiration data data\_r <- calculate\_r(data = data\_env, leaf\_Q10\_day = 2.015, #Atkin & Tjoelker 2003, 25C acclimated leaf\_Q10\_night = 2.015, root\_Q10 = 2.015, shoot\_Q10 = 2.015, leaf\_k25\_day = 2.78 \* 0.7, #Assumption from Stinziano et al. 2019 leaf\_k25\_night = 2.78, #Resp from control in Stinziano & Way 2017 root\_k25 = 0.0095,#Weger and Guy 1991 shoot\_k25 = 0.0095,#Assume stem and root respiration rates are similar leaf\_area = leaf\_area\_1, root\_mass = root\_mass\_1, shoot\_mass = stem\_mass\_1) data\_r <- data\_r[, 1:7] #remove temperature column - poses issue later for(i in 1:nrow(pars\_v)){ data\_p <- calculate\_p(G25 = 42.75, #Bernacchi et al. 2001 G\_Ea = 37.830, #Bernacchi et al. 2001 Km25 = 718.40, #Bernacchi et al. 2001 Km\_Ea = 65.50828, #Bernacchi et al. 2001 Temp = data\_env\(Temp, Vcmax25\_m = pars\_v\)k25\_o[i], Vcmax\_dS\_m = pars\_v\(dS\_o[i], Vcmax\_Hd\_m = pars\_v\)Hd\_o[i], Vcmax\_Ea\_m = pars\_v\(Ea\_o[i], Vcmax25\_n = pars\_v\)k25[i], Vcmax\_dS\_n = pars\_v\(dS[i], Vcmax\_Hd\_n = pars\_v\)Hd[i], Vcmax\_Ea\_n = pars\_v\(Ea[i], Jmax25\_m = pars\_j\)k25\_o[i], Jmax\_dS\_m = pars\_j\(dS\_o[i], Jmax\_Hd\_m = pars\_j\)Hd\_o[i], Jmax\_Ea\_m = pars\_j\(Ea\_o[i], Jmax25\_n = pars\_j\)k25[i], Jmax\_dS\_n = pars\_j\(dS[i], Jmax\_Hd\_n = pars\_j\)Hd[i], Jmax\_Ea\_n = pars\_j$Ea[i]) data\_p <- data\_p[, 1:6] #Removes temp column to preclude issues later data\_phys[[i]] <- cbind(data\_r, data\_p) }

#Model with Medlyn equation modelled <- list(NULL) for(i in 1:length(data\_phys)){ modelled[[i]] <- calculate\_photo(data\_phys = data\_phys[[i]], data\_env = data\_env, varnames = list(GammaStar = “GammaStar”, Jmax = “Jmax\_medlyn”, Km = “Km”, leaf\_respiration = “leaf\_respiration”, root\_respiration = “root\_respiration”, shoot\_respiration = “shoot\_respiration”, total\_respiration\_plant = “total\_respiration\_plant”, Ca = “Ca”, Qin = “Qin”, VPD = “VPD”, Temp = “Temp”, Vcmax = “Vcmax\_medlyn”), leaf\_area = leaf\_area\_1, #phi = 0.08, #Campbell & Norman 1998 phi = 0.85, #CLM 5.0 time\_step = 3600) #3600 s/hr }

for(i in 1:length(modelled)){ modelled[[i]][[1]]$ID <- “Medlyn” }

ge <- list(NULL) for(i in 1:length(modelled)){ ge[[i]] <- modelled[[i]][[1]] }

ge <- do.call(“rbind”, ge)

dailyC <- list(NULL) for(i in 1:length(modelled)){ dailyC[[i]] <- modelled[[i]][[1]] %>% group\_by(DOY, Location) %>% summarize(daily\_plant\_C = sum(A\_net\_plant \* 3600) \* 12.01 / 1000000, #12.01 g/mol, umol to mol daily\_plant\_photosynthesis = sum(A\_gross\_plant \* 3600) \* 12.01 / 1000000, daily\_plant\_respiration = sum(total\_respiration\_plant \* 3600) \* 12.01 / 1000000) dailyC[[i]]$ID <- “Medlyn” } dailyC <- do.call(“rbind”, dailyC)

#Model with rederivation modelled2 <- list(NULL) for(i in 1:length(data\_phys)){ modelled2[[i]] <- calculate\_photo(data\_phys = data\_phys[[i]], data\_env = data\_env, varnames = list(GammaStar = “GammaStar”, Jmax = “Jmax\_new”, Km = “Km”, leaf\_respiration = “leaf\_respiration”, root\_respiration = “root\_respiration”, shoot\_respiration = “shoot\_respiration”, total\_respiration\_plant = “total\_respiration\_plant”, Ca = “Ca”, Qin = “Qin”, VPD = “VPD”, Temp = “Temp”, Vcmax = “Vcmax\_new”), leaf\_area = leaf\_area\_1, phi = 0.85, time\_step = 3600) }

for(i in 1:length(modelled2)){ modelled2[[i]][[1]]$ID <- “New” }

ge2 <- list(NULL) for(i in 1:length(modelled2)){ ge2[[i]] <- modelled2[[i]][[1]] }

ge2 <- do.call(“rbind”, ge2)

dailyC2 <- list(NULL) for(i in 1:length(modelled2)){ dailyC2[[i]] <- modelled2[[i]][[1]] %>% group\_by(DOY, Location) %>% summarize(daily\_plant\_C = sum(A\_net\_plant \* 3600) \* 12.01 / 1000000, daily\_plant\_photosynthesis = sum(A\_gross\_plant \* 3600) \* 12.01 / 1000000, daily\_plant\_respiration = sum(total\_respiration\_plant \* 3600) \* 12.01 / 1000000) dailyC2[[i]]$ID <- “New” } dailyC2 <- do.call(“rbind”, dailyC2)

week\_1\_carbon\_Hdfit <- rbind(dailyC, dailyC2) week\_1\_ge <- rbind(ge, ge2)

boxplot(daily\_plant\_C ~ ID, data = week\_1\_carbon\_Hdfit) summary(lm(week\_1\_carbon\_Hdfit[week\_1\_carbon\_Hdfit$ID == “Medlyn”,]\(daily\_plant\_C ~ week\_1\_carbon\_Hdfit[week\_1\_carbon\_Hdfit\)ID == “New”,]\(daily\_plant\_C-1)) #0.1% underestimation lm(week\_1\_carbon\_Hdfit[week\_1\_carbon\_Hdfit\)ID == “Medlyn”,]\(daily\_plant\_C ~ week\_1\_carbon\_Hdfit[week\_1\_carbon\_Hdfit\)ID == “New”,]$daily\_plant\_C)

```
#Statistics and graphical output for Week 1 (Hd Fit)
#Remove octothorpe below to run chunk
#```{r include = FALSE}
# Additional column for daily plant A/R
week_1_carbon_Hdfit$daily_plant_AR <- (week_1_carbon_Hdfit$daily_plant_photosynthesis/week_1_carbon_Hdfit$daily_plant_respiration)
week_1_carbon_Hdfit$daily_plant_AR

#Stats
summary(lm(week_1_carbon_Hdfit[week_1_carbon_Hdfit$ID == "Medlyn",]$daily_plant_C ~ week_1_carbon_Hdfit[week_1_carbon_Hdfit$ID == "New",]$daily_plant_C))

summary(lm(week_1_carbon_Hdfit[week_1_carbon_Hdfit$ID == "Medlyn",]$daily_plant_photosynthesis ~ week_1_carbon_Hdfit[week_1_carbon_Hdfit$ID == "New",]$daily_plant_photosynthesis))

summary(lm(week_1_carbon_Hdfit[week_1_carbon_Hdfit$ID == "Medlyn",]$daily_plant_AR ~ week_1_carbon_Hdfit[week_1_carbon_Hdfit$ID == "New",]$daily_plant_AR))

#Basic Graphs
plot(week_1_carbon_Hdfit[week_1_carbon_Hdfit$ID == "New",]$daily_plant_C ~ week_1_carbon_Hdfit[week_1_carbon_Hdfit$ID == "Medlyn",]$daily_plant_C,
  xlab="New Total Daily C Gain", ylab="Old Total Daily C Gain", main="a) Week 1 using Hd Fit")
abline(lm(week_1_carbon_Hdfit[week_1_carbon_Hdfit$ID == "New",]$daily_plant_C ~ week_1_carbon_Hdfit[week_1_carbon_Hdfit$ID == "Medlyn",]$daily_plant_C), col="blue")

plot(week_1_carbon_Hdfit[week_1_carbon_Hdfit$ID == "New",]$daily_plant_photosynthesis ~ week_1_carbon_Hdfit[week_1_carbon_Hdfit$ID == "Medlyn",]$daily_plant_photosynthesis,
  xlab="New Total Daily Photosynthesis", ylab="Old Total Daily Photosynthesis", main="c) Week 1 using Hd Fit")
abline(lm(week_1_carbon_Hdfit[week_1_carbon_Hdfit$ID == "New",]$daily_plant_photosynthesis ~ week_1_carbon_Hdfit[week_1_carbon_Hdfit$ID == "Medlyn",]$daily_plant_photosynthesis), col="blue")

plot(week_1_carbon_Hdfit[week_1_carbon_Hdfit$ID == "New",]$daily_plant_AR ~ week_1_carbon_Hdfit[week_1_carbon_Hdfit$ID == "Medlyn",]$daily_plant_AR,
  xlab="New Total Daily A/R", ylab="Old Total Daily A/R", main="e) Week 1 using Hd Fit")
abline(lm(week_1_carbon_Hdfit[week_1_carbon_Hdfit$ID == "New",]$daily_plant_AR ~ week_1_carbon_Hdfit[week_1_carbon_Hdfit$ID == "Medlyn",]$daily_plant_AR), col="blue")
```

#Week 12 modeling - high root/leaf ratio #Remove octothorpe below to run chunk #```{r include = FALSE} data <- read.csv(system.file(“extdata”, “env\_data.csv”, package = “arrhenius.comparison”), stringsAsFactors = FALSE) pars\_v <- read.csv(system.file(“extdata”, “fits\_v\_3.csv”, package = “arrhenius.comparison”), stringsAsFactors = FALSE) pars\_j <- read.csv(system.file(“extdata”, “fits\_j\_3.csv”, package = “arrhenius.comparison”), stringsAsFactors = FALSE) pars\_v <- pars\_v[pars\_v$Hd != 200,] pars\_j <- pars\_v[pars\_j$Hd != 200,] data\_phys <- list(NULL) #Basically, we need a solid set of respiration data data\_r <- calculate\_r(data = data, leaf\_Q10\_day = 2.015, leaf\_Q10\_night = 2.015, root\_Q10 = 2.015, shoot\_Q10 = 2.015, leaf\_k25\_day = 2.78 \* 0.7, leaf\_k25\_night = 2.78, root\_k25 = 0.0095,#Weger and Guy 1991 shoot\_k25 = 0.0095,#Assume stem and root respiration rates are similar leaf\_area = leaf\_area\_12, root\_mass = root\_mass\_12, shoot\_mass = stem\_mass\_12) data\_r <- data\_r[, 1:7] #remove temperature column - poses issue later for(i in 1:nrow(pars\_v)){ data\_p <- calculate\_p(G25 = 42.75, G\_Ea = 37.830, Km25 = 718.40, Km\_Ea = 65.50828, Temp = data\(Temp, Vcmax25\_m = pars\_v\)k25\_o[i], Vcmax\_dS\_m = pars\_v\(dS\_o[i], Vcmax\_Hd\_m = pars\_v\)Hd\_o[i], Vcmax\_Ea\_m = pars\_v\(Ea\_o[i], Vcmax25\_n = pars\_v\)k25[i], Vcmax\_dS\_n = pars\_v\(dS[i], Vcmax\_Hd\_n = pars\_v\)Hd[i], Vcmax\_Ea\_n = pars\_v\(Ea[i], Jmax25\_m = pars\_j\)k25\_o[i], Jmax\_dS\_m = pars\_j\(dS\_o[i], Jmax\_Hd\_m = pars\_j\)Hd\_o[i], Jmax\_Ea\_m = pars\_j\(Ea\_o[i], Jmax25\_n = pars\_j\)k25[i], Jmax\_dS\_n = pars\_j\(dS[i], Jmax\_Hd\_n = pars\_j\)Hd[i], Jmax\_Ea\_n = pars\_j$Ea[i]) data\_p <- data\_p[, 1:6] #Removes temp column to preclude issues later data\_phys[[i]] <- cbind(data\_r, data\_p) data\_phys }

modelled <- list(NULL) for(i in 1:length(data\_phys)){ modelled[[i]] <- calculate\_photo(data\_phys = data\_phys[[i]], data\_env = data, varnames = list(GammaStar = “GammaStar”, Jmax = “Jmax\_medlyn”, Km = “Km”, leaf\_respiration = “leaf\_respiration”, root\_respiration = “root\_respiration”, shoot\_respiration = “shoot\_respiration”, total\_respiration\_plant = “total\_respiration\_plant”, Ca = “Ca”, Qin = “Qin”, VPD = “VPD”, Temp = “Temp”, Vcmax = “Vcmax\_medlyn”), leaf\_area = leaf\_area\_12, phi = 0.85, time\_step = 3600) }

#Note, we will need a for loop that looks like this: for(i in 1:length(modelled)){ modelled[[i]][[1]]$ID <- “Medlyn” }

ge <- list(NULL) for(i in 1:length(modelled)){ ge[[i]] <- modelled[[i]][[1]] }

ge <- do.call(“rbind”, ge)

dailyC <- list(NULL) for(i in 1:length(modelled)){ dailyC[[i]] <- modelled[[i]][[1]] %>% group\_by(DOY, Location) %>% summarize(daily\_plant\_C = sum(A\_net\_plant \* 3600) \* 12.01 / 1000000, daily\_plant\_photosynthesis = sum(A\_gross\_plant \* 3600) \* 12.01 / 1000000, daily\_plant\_respiration = sum(total\_respiration\_plant \* 3600) \* 12.01 / 1000000) dailyC[[i]]$ID <- “Medlyn” } dailyC <- do.call(“rbind”, dailyC)

modelled2 <- list(NULL) for(i in 1:length(data\_phys)){ modelled2[[i]] <- calculate\_photo(data\_phys = data\_phys[[i]], data\_env = data, varnames = list(GammaStar = “GammaStar”, Jmax = “Jmax\_new”, Km = “Km”, leaf\_respiration = “leaf\_respiration”, root\_respiration = “root\_respiration”, shoot\_respiration = “shoot\_respiration”, total\_respiration\_plant = “total\_respiration\_plant”, Ca = “Ca”, Qin = “Qin”, VPD = “VPD”, Temp = “Temp”, Vcmax = “Vcmax\_new”), leaf\_area = leaf\_area\_12, phi = 0.85, time\_step = 3600) } #Note, we will need a for loop that looks like this: for(i in 1:length(modelled2)){ modelled2[[i]][[1]]$ID <- “New” }

ge2 <- list(NULL) for(i in 1:length(modelled2)){ ge2[[i]] <- modelled2[[i]][[1]] }

ge2 <- do.call(“rbind”, ge2)

dailyC2 <- list(NULL) for(i in 1:length(modelled2)){ dailyC2[[i]] <- modelled2[[i]][[1]] %>% group\_by(DOY, Location) %>% summarize(daily\_plant\_C = sum(A\_net\_plant \* 3600) \* 12.01 / 1000000, daily\_plant\_photosynthesis = sum(A\_gross\_plant \* 3600) \* 12.01 / 1000000, daily\_plant\_respiration = sum(total\_respiration\_plant \* 3600) \* 12.01 / 1000000) dailyC2[[i]]$ID <- “New” } dailyC2 <- do.call(“rbind”, dailyC2)

week\_12\_carbon\_Hdfit <- rbind(dailyC, dailyC2) week\_12\_ge <- rbind(ge, ge2) boxplot(daily\_plant\_C ~ ID, data = week\_12\_carbon\_Hdfit) summary(lm(week\_12\_carbon\_Hdfit[week\_12\_carbon\_Hdfit$ID == “Medlyn”,]\(daily\_plant\_C ~ week\_12\_carbon\_Hdfit[week\_12\_carbon\_Hdfit\)ID == “New”,]$daily\_plant\_C-1)) #0.1% underestimation

```
#Statistics and graphical output for Week 12 (Hd fit)
#Remove octothorpe below to run chunk
#```{r include = FALSE}
# Additional column for daily plant A/R
week_12_carbon_Hdfit$daily_plant_AR <- (week_12_carbon_Hdfit$daily_plant_photosynthesis/week_12_carbon_Hdfit$daily_plant_respiration)
week_12_carbon_Hdfit$daily_plant_AR

#Stats
summary(lm(week_12_carbon_Hdfit[week_12_carbon_Hdfit$ID == "Medlyn",]$daily_plant_C ~ week_12_carbon_Hdfit[week_12_carbon_Hdfit$ID == "New",]$daily_plant_C))

summary(lm(week_12_carbon_Hdfit[week_12_carbon_Hdfit$ID == "Medlyn",]$daily_plant_photosynthesis ~ week_12_carbon_Hdfit[week_12_carbon_Hdfit$ID == "New",]$daily_plant_photosynthesis))

summary(lm(week_12_carbon_Hdfit[week_12_carbon_Hdfit$ID == "Medlyn",]$daily_plant_AR ~ week_12_carbon_Hdfit[week_12_carbon_Hdfit$ID == "New",]$daily_plant_AR))

#Graphs
plot(week_12_carbon_Hdfit[week_12_carbon_Hdfit$ID == "New",]$daily_plant_C ~ week_12_carbon_Hdfit[week_12_carbon_Hdfit$ID == "Medlyn",]$daily_plant_C,
  xlab="New Total Daily C Gain", ylab="Old Total Daily C Gain", main="b) Week 12 using Hd Fit")
abline(lm(week_12_carbon_Hdfit[week_12_carbon_Hdfit$ID == "New",]$daily_plant_C ~ week_12_carbon_Hdfit[week_12_carbon_Hdfit$ID == "Medlyn",]$daily_plant_C), col="blue")

plot(week_12_carbon_Hdfit[week_12_carbon_Hdfit$ID == "New",]$daily_plant_photosynthesis ~ week_12_carbon_Hdfit[week_12_carbon_Hdfit$ID == "Medlyn",]$daily_plant_photosynthesis,
  xlab="New Total Daily Photosynthesis", ylab="Old Total Daily Photosynthesis", main="d) Week 12 using Hd Fit")
abline(lm(week_12_carbon_Hdfit[week_12_carbon_Hdfit$ID == "New",]$daily_plant_photosynthesis ~ week_12_carbon_Hdfit[week_12_carbon_Hdfit$ID == "Medlyn",]$daily_plant_photosynthesis), col="blue")

plot(week_12_carbon_Hdfit[week_12_carbon_Hdfit$ID == "New",]$daily_plant_AR ~ week_12_carbon_Hdfit[week_12_carbon_Hdfit$ID == "Medlyn",]$daily_plant_AR,
  xlab="New Total Daily A/R", ylab="Old Total Daily A/R", main="f) Week 12 using Hd Fit")
abline(lm(week_12_carbon_Hdfit[week_12_carbon_Hdfit$ID == "New",]$daily_plant_AR ~ week_12_carbon_Hdfit[week_12_carbon_Hdfit$ID == "Medlyn",]$daily_plant_AR), col="blue")
```

Merge both datasets and output #Remove octothorpe below to run chunk #`{r include = FALSE} carbon <- rbind(week_1_carbon, week_12_carbon) gasex <- rbind(week_1_ge, week_12_ge) write.csv(carbon, "carbon_HdFit.csv") write.csv(gasex, "gasex_HdFit.csv")` #Remove octothorpe below to run chunk #```{r include = FALSE} #More complex ggplot graphs FigData <- read.csv(system.file(“extdata”, “FigModelingData.csv”, package = “arrhenius.comparison”), stringsAsFactors = FALSE)

#Daily C Gain Graphs W1\_Hd200\_CGain <- ggplot(data = FigData, aes(x = plant\_C\_.medlyn\_W1\_fix, y = plant\_C\_new\_W1\_fix)) + ggtitle(label = expression(“b)”_“Low”_“R”_“With”_“Fixed”~H[d])) + geom\_point(size = 3, alpha = 0.7)+ geom\_abline(slope = 1, intercept = 0, size = 1, colour = “Black”) + geom\_abline(slope = 0.978097, intercept = 0.001426, size = 2, linetype = “dashed”, colour = “Gray45”)+ theme\_bw() + theme(panel.grid = element\_blank())+ theme(text = element\_text(size = 12, colour = “Black”), axis.line=element\_line(size=1), axis.ticks=element\_line(size=1.5), axis.ticks.length=unit(1.5, “mm”), rect=element\_rect(size=2))+ xlim(-0.3,0.3)+ ylim(-0.3,0.3)+ labs(x=expression(‘Old’_‘Total’_‘C’_‘Gain’_“(gC”~“plant”^{-1}~“day”^{-1}*“)”), y=expression(‘New’_‘Total’_‘C’_‘Gain’_“(gC”~“plant”^{-1}~“day”^{-1}*“)”)) W1\_Hd200\_CGain

W12\_Hd200\_CGain <- ggplot(data = FigData, aes(x = plant\_C\_medlyn\_W12\_fix, y = plant\_C\_new\_W12\_fix)) + ggtitle(label = expression(“d)”_“High”_“R”_“With”_“Fixed”~H[d])) + geom\_point(size = 3, alpha = 0.7)+ geom\_abline(slope = 1, intercept = 0, size = 1, colour = “Black”) + geom\_abline(slope = 0.979928, intercept = 0.001377, size = 2, linetype = “dashed”, colour = “Gray45”)+ theme\_bw() + theme(panel.grid = element\_blank())+ theme(text = element\_text(size = 12, colour = “Black”), axis.line=element\_line(size=1), axis.ticks=element\_line(size=1.5), axis.ticks.length=unit(1.5, “mm”), rect=element\_rect(size=2))+ xlim(-0.3,0.3)+ ylim(-0.3,0.3)+ labs(x=expression(‘Old’_‘Total’_‘C’_‘Gain’_“(gC”~“plant”^{-1}~“day”^{-1}*“)”), y=expression(‘New’_‘Total’_‘C’_‘Gain’_“(gC”~“plant”^{-1}~“day”^{-1}*“)”)) W12\_Hd200\_CGain

names(FigData) W1\_HdVar\_CGain <- ggplot(data = FigData, aes(x = plant\_C\_.medlyn\_W1\_var, y = plant\_C\_.new\_W1\_var)) + ggtitle(label = expression(“a)”_“Low”_“R”_“With”_“Variable”~H[d])) + geom\_point(size = 3, alpha = 0.7)+ geom\_abline(slope = 1, intercept = 0, size = 1, colour = “Black”) + geom\_abline(slope = 0.983546, intercept = 0.001233, size = 2, linetype = “dashed”, colour = “Gray45”)+ theme\_bw() + theme(panel.grid = element\_blank())+ theme(text = element\_text(size = 12, colour = “Black”), axis.line=element\_line(size=1), axis.ticks=element\_line(size=1.5), axis.ticks.length=unit(1.5, “mm”), rect=element\_rect(size=2))+ xlim(-0.3,0.3)+ ylim(-0.3,0.3)+ labs(x=expression(‘Old’_‘Total’_‘C’_‘Gain’_“(gC”~“plant”^{-1}~“day”^{-1}*“)”), y=expression(‘New’_‘Total’_‘C’_‘Gain’_“(gC”~“plant”^{-1}~“day”^{-1}*“)”)) W1\_HdVar\_CGain

W12\_HdVar\_CGain <- ggplot(data = FigData, aes(x = plant\_C\_.medlyn\_W12\_var, y = plant\_C\_.new\_W12\_var)) + ggtitle(label = expression(“c)”_“High”_“R”_“With”_“Variable”~H[d])) + geom\_point(size = 3, alpha = 0.7)+ geom\_abline(slope = 1, intercept = 0, size = 1, colour = “Black”) + geom\_abline(slope = 0.985405, intercept = 0.001282, size = 2, linetype = “dashed”, colour = “Gray45”)+ theme\_bw() + theme(panel.grid = element\_blank())+ theme(text = element\_text(size = 12, colour = “Black”), axis.line=element\_line(size=1), axis.ticks=element\_line(size=1.5), axis.ticks.length=unit(1.5, “mm”), rect=element\_rect(size=2))+ xlim(-0.3,0.3)+ ylim(-0.3,0.3)+ labs(x=expression(‘Old’_‘Total’_‘C’_‘Gain’_“(gC”~“plant”^{-1}~“day”^{-1}*“)”), y=expression(‘New’_‘Total’_‘C’_‘Gain’_“(gC”~“plant”^{-1}~“day”^{-1}*“)”)) W12\_HdVar\_CGain

library(gridExtra) jpeg(“Figure 6.jpeg”, width = 7, height = 7, units = “in”, res = 600) grid.arrange(W1\_HdVar\_CGain, W1\_Hd200\_CGain, W12\_HdVar\_CGain, W12\_Hd200\_CGain, ncol = 2, nrow = 2) dev.off()

#### Daily Photosynthesis Across Models

names(FigData) W1\_Hd200\_Photo <- ggplot(data = FigData, aes(x = photo\_medlyn\_W1\_fix, y = photo\_new\_W1\_fix)) + ggtitle(label = expression(“b)”_“Low”_“R”_“With”_“Fixed”~H[d])) + geom\_point(size = 3, alpha = 0.7)+ geom\_abline(slope = 1, intercept = 0, size = 1, colour = “Black”) + geom\_abline(slope = 0.978097, intercept = 0.001426, size = 2, linetype = “dashed”, colour = “Gray45”)+ theme\_bw() + xlim(-0.15,0.4)+ ylim(-0.15,0.4)+ theme(panel.grid = element\_blank())+ theme(text = element\_text(size = 12, colour = “Black”), axis.line=element\_line(size=1), axis.ticks=element\_line(size=1.5), axis.ticks.length=unit(1.5, “mm”), rect=element\_rect(size=2))+ labs(x=expression(‘Old’_‘Total’_‘Daily’_‘A’_“(gC”~“plant”^{-1}~“day”^{-1}*“)”), y=expression(‘New’_‘Total’_‘Daily’_‘A’_“(gC”~“plant”^{-1}~“day”^{-1}*“)”)) W1\_Hd200\_Photo

W12\_Hd200\_Photo <- ggplot(data = FigData, aes(x = photo\_medlyn\_W12\_fix, y = photo\_new\_W12\_fix)) + ggtitle(label = expression(“d)”_“High”_“R”_“With”_“Fixed”~H[d])) + geom\_point(size = 3, alpha = 0.7)+ geom\_abline(slope = 1, intercept = 0, size = 1, colour = “Black”) + geom\_abline(slope = 0.979928, intercept = 0.001377, size = 2, linetype = “dashed”, colour = “Gray45”)+ theme\_bw() + theme(panel.grid = element\_blank())+ theme(text = element\_text(size = 12, colour = “Black”), axis.line=element\_line(size=1), axis.ticks=element\_line(size=1.5), axis.ticks.length=unit(1.5, “mm”), rect=element\_rect(size=2))+ xlim(-0.15,0.4)+ ylim(-0.15,0.4)+ labs(x=expression(‘Old’_‘Total’_‘Daily’_‘A’_“(gC”~“plant”^{-1}~“day”^{-1}*“)”), y=expression(‘New’_‘Total’_‘Daily’_‘A’_“(gC”~“plant”^{-1}~“day”^{-1}*“)”)) W12\_Hd200\_Photo

names(FigData) W1\_HdVar\_Photo <- ggplot(data = FigData, aes(x = photo\_medlyn\_W1\_var, y = photo\_new\_W1\_var)) + ggtitle(label = expression(“a)”_“Low”_“R”_“With”_“Variable”~H[d])) + geom\_point(size = 3, alpha = 0.7)+ geom\_abline(slope = 1, intercept = 0, size = 1, colour = “Black”) + geom\_abline(slope = 0.983546, intercept = 0.001233, size = 2, linetype = “dashed”, colour = “Gray45”)+ theme\_bw() + theme(panel.grid = element\_blank())+ theme(text = element\_text(size = 12, colour = “Black”), axis.line=element\_line(size=1), axis.ticks=element\_line(size=1.5), axis.ticks.length=unit(1.5, “mm”), rect=element\_rect(size=2))+ xlim(-0.15,0.4)+ ylim(-0.15,0.4)+ labs(x=expression(‘Old’_‘Total’_‘Daily’_‘A’_“(gC”~“plant”^{-1}~“day”^{-1}*“)”), y=expression(‘New’_‘Total’_‘Daily’_‘A’_“(gC”~“plant”^{-1}~“day”^{-1}*“)”)) W1\_HdVar\_Photo

W12\_HdVar\_Photo <- ggplot(data = FigData, aes(x = photo\_medlyn\_W12\_var, y = photo\_new\_W12\_var)) + ggtitle(label = expression(“c)”_“High”_“R”_“With”_“Variable”~H[d])) + geom\_point(size = 3, alpha = 0.7)+ geom\_abline(slope = 1, intercept = 0, size = 1, colour = “Black”) + geom\_abline(slope = 0.985405, intercept = 0.001282, size = 2, linetype = “dashed”, colour = “Gray45”)+ theme\_bw() + theme(panel.grid = element\_blank())+ theme(text = element\_text(size = 12, colour = “Black”), axis.line=element\_line(size=1), axis.ticks=element\_line(size=1.5), axis.ticks.length=unit(1.5, “mm”), rect=element\_rect(size=2))+ xlim(-0.15,0.4)+ ylim(-0.15,0.4)+ labs(x=expression(‘Old’_‘Total’_‘Daily’_‘A’_“(gC”~“plant”^{-1}~“day”^{-1}*“)”), y=expression(‘New’_‘Total’_‘Daily’_‘A’_“(gC”~“plant”^{-1}~“day”^{-1}*“)”)) W12\_HdVar\_Photo

library(gridExtra) jpeg(“Figure 4.jpeg”, width = 7, height = 7, units = “in”, res = 600) grid.arrange(W1\_HdVar\_Photo, W1\_Hd200\_Photo, W12\_HdVar\_Photo, W12\_Hd200\_Photo, ncol = 2, nrow = 2) dev.off()

#### Daily A:R

names(FigData) W1\_Hd200\_AR <- ggplot(data = FigData, aes(x = AR\_medlyn\_W1\_fix, y = AR\_new\_W1\_fix)) + ggtitle(label = expression(“b)”_“Low”_“R”_“With”_“Fixed”~H[d])) + geom\_point(size = 3, alpha = 0.7)+ geom\_abline(slope = 1, intercept = 0, size = 1, colour = “Black”) + geom\_abline(slope = 0.978097, intercept = 0.001426, size = 2, linetype = “dashed”, colour = “Gray45”)+ theme\_bw() + xlim(-0.5,9.5)+ ylim(-0.5,9.5)+ theme(panel.grid = element\_blank())+ theme(text = element\_text(size = 12, colour = “Black”), axis.line=element\_line(size=1), axis.ticks=element\_line(size=1.5), axis.ticks.length=unit(1.5, “mm”), rect=element\_rect(size=2))+ labs(x=expression(‘Old’_‘Total’_‘Daily’~‘A/R’), y=expression(‘New’_‘Total’_‘Daily’~‘A/R’)) W1\_Hd200\_AR

W12\_Hd200\_AR <- ggplot(data = FigData, aes(x = AR\_medlyn\_W12\_fix, y = AR\_new\_W12\_fix)) + ggtitle(label = expression(“d)”_“High”_“R”_“With”_“Fixed”~H[d])) + geom\_point(size = 3, alpha = 0.7)+ geom\_abline(slope = 1, intercept = 0, size = 1, colour = “Black”) + geom\_abline(slope = 0.979928, intercept = 0.001377, size = 2, linetype = “dashed”, colour = “Gray45”)+ theme\_bw() + theme(panel.grid = element\_blank())+ theme(text = element\_text(size = 12, colour = “Black”), axis.line=element\_line(size=1), axis.ticks=element\_line(size=1.5), axis.ticks.length=unit(1.5, “mm”), rect=element\_rect(size=2))+ xlim(-0.5,9.5)+ ylim(-0.5,9.5)+ labs(x=expression(‘Old’_‘Total’_‘Daily’~‘A/R’), y=expression(‘New’_‘Total’_‘Daily’~‘A/R’)) W12\_Hd200\_AR

W1\_HdVar\_AR <- ggplot(data = FigData, aes(x = AR\_medlyn\_W1\_var, y = AR\_new\_W1\_var)) + ggtitle(label = expression(“a)”_“Low”_“R”_“With”_“Variable”~H[d])) + geom\_point(size = 3, alpha = 0.7)+ geom\_abline(slope = 1, intercept = 0, size = 1, colour = “Black”) + geom\_abline(slope = 0.983546, intercept = 0.001233, size = 2, linetype = “dashed”, colour = “Gray45”)+ theme\_bw() + theme(panel.grid = element\_blank())+ theme(text = element\_text(size = 12, colour = “Black”), axis.line=element\_line(size=1), axis.ticks=element\_line(size=1.5), axis.ticks.length=unit(1.5, “mm”), rect=element\_rect(size=2))+ xlim(-0.5,9.5)+ ylim(-0.5,9.5)+ labs(x=expression(‘Old’_‘Total’_‘Daily’~‘A/R’), y=expression(‘New’_‘Total’_‘Daily’~‘A/R’)) W1\_HdVar\_AR

W12\_HdVar\_AR <- ggplot(data = FigData, aes(x = AR\_medlyn\_W12\_var, y = AR\_new\_W12\_var)) + ggtitle(label = expression(“c)”_“High”_“R”_“With”_“Variable”~H[d])) + geom\_point(size = 3, alpha = 0.7)+ geom\_abline(slope = 1, intercept = 0, size = 1, colour = “Black”) + geom\_abline(slope = 0.985405, intercept = 0.001282, size = 2, linetype = “dashed”, colour = “Gray45”)+ theme\_bw() + theme(panel.grid = element\_blank())+ theme(text = element\_text(size = 12, colour = “Black”), axis.line=element\_line(size=1), axis.ticks=element\_line(size=1.5), axis.ticks.length=unit(1.5, “mm”), rect=element\_rect(size=2))+ xlim(-0.5,9.5)+ ylim(-0.5,9.5)+ labs(x=expression(‘Old’_‘Total’_‘Daily’~‘A/R’), y=expression(‘New’_‘Total’_‘Daily’~‘A/R’)) W12\_HdVar\_AR

library(gridExtra) jpeg(“Figure 5.jpeg”, width = 7.5, height = 7.5, units = “in”, res = 600) grid.arrange(W1\_HdVar\_AR, W1\_Hd200\_AR, W12\_HdVar\_AR, W12\_Hd200\_AR, ncol = 2, nrow = 2) dev.off()  
<<<<<<< HEAD

#### Compound interest comparsions

#Initial biomass of 1g #180 day growing season #Lowest % difference = 1.5% for week 12 fitted Medlyn = 0.0563 gC plant-1 day-1 New = 0.0558 gC plant-1 day-1

#Daily Interest FV <- PV\*((1+i)^n

FV = (1)\*(1+(0.0558))^180 FV

Med <- 0.0558*180 = 10.044 g + 1 g = 11.044 g New <- 0.0563*180 = 10.134 g + 1 g = 11.134 g Diff <- <1% (0.815%)

#Compound Interest P=1 n=180 R1=(0.0558) R2=(0.0563)

A1=(P*(1+(R1/1))^n) A2=(P*(1+(R2/1))^n) A1 A2

Diff=((A2-A1)/((A1+A2)/2))\*100 Diff

======= ``` >>>>>>> 1a689f9231e4e8257d522c97bfa2792ed2884f6d
