## Supplementary material for "A derivation error that affects carbon balance models exists in the current implementation of the modified Arrhenius function": R package for analysis: Murphy_Stinziano_Reviewer_Response.docx (from another version of this preprint: version 2, 2020-04-23)

**Response to reviewers**

We have addressed referee reports from our *Global Change Biology* submission (ID: GCB-20-0170) below and believe that our manuscript has been strengthened by addressing the reviewer comments. Our responses are in bold, and we have attached a version of the manuscript with tracked changes for review only.

**Reviewer Comments from GCB Submission**

*Reviewer: 1*
Comments to the Author
This manuscript describes a comparison of modeled carbon balance using a modern version of a temperature response function originally derived in 1942 with the original version of that function. The authors noticed that the modern version omits a direct proportionality to temperature itself (absolute temperature, in kelvins). The reason for the omission is unclear; perhaps it was a mathematical error in earlier works applying the 1942 expression. The authors show that if the original version of the response function is used instead of the modern version (the one with the apparent error), predictions of carbon balance are substantially affected.

I appreciate the attention to detail that brought the authors to notice this odd omission, which has propagated through the recent literature involving temperature responses of parameters in plant ecophysiology. In principle, the approach is also broadly sound and valuable. However, I have some significant concerns regarding the relevance of the comparison, and consequently the value of the authors' interpretation of the result.

Few, if any, modern ecophysiologists, and particularly those engaged in modeling at large scales, interpret these temperature response functions and their internal parameters as essentially mechanistic. That is, the internal parameters are *never* estimated on the basis of their original mechanistic interpretation in terms of biochemistry. Instead, the equations are always fitted empirically to available temperature response data. This is perhaps most strongly illustrated by the fact that different functions have emerged as the preferred ones for simulating T responses of various photosynthetic and respiratory parameters. Those choices have generally not been based on biochemical principles, but rather, on how well any given function fitted the data.

**We appreciate the reviewer’s comments here, however we did not make these points nor argue for them in our manuscript.**

Therefore, the figure of merit for "temperature response functions" isn't whether they are faithful to their original derivations. What matters in practice is how well they fit the data. It is in this respect that the current ms falls short: there is literally no attention given to, nor any information provided about, whether the "old" or "new" equation leads to more *accurate* predictions of either T responses of variables, narrowly, or of carbon balance more broadly. In my mind, that is the one and only important question from the standpoint of modeling carbon balance. Unfortunately the current ms (as written) doesn't address this point at all. Instead it focuses on differences in the fitted internal parameters (delta-H, etc).

**We thank the reviewer for their comments and now address the “accuracy” of the two equations (Fig. 2, lines 211 – 216). Philosophically speaking, the current implementation of the modified Arrhenius function is logically in error. Therefore, even though our new analysis shows that both equations have nearly identical performance, the logical error means that the new derivation should be used instead. We do not believe that an equation with logical errors should be used for fitting and modeling data.**

To be clear, I fully understand the value of mechanistic models as opposed to empirical ones. I recognize that the internal parameters might have some conserved meaning. But these models aren't really applied in this way. For example, people don't (to my knowledge) predict acclimation or adaptation of T response curves on the basis of conservative principles involving these internal parameters; instead they do experiments and re-fit the equations empirically. It's a bit like the difference between how gamma-star is estimated empirically from gas exchange experiments instead of being estimated using experimental values for its component parameters.

**We no longer argue for a specific move away from the modified Arrhenius equation and therefore no longer need to address this point.**

There are some important omissions from the ms, such as the temperature range of the data to which the two equations were fitted vs applied in the carbon balance modeling, and information about the experiments that produced those data. It would be very helpful also to see the actual curves, both in generic terms and as specifically fitted to the data. Perhaps also as fitted to one another, to get a sense of how different they really are in principle.

**We now show example temperature responses of V_cmax_ and J_max_ with cases where the curves are virtually identical and where they are different (Fig. 2c-f). We also now outline the temperature ranges of the curve fits (lines 153-154) and the environmental data (lines 191-193) used to drive the model, which are fairly comparable: -0.1 – 42.2 °C for environmental data, 1 – 50 °C for the curve fits. All of the information on the experiments from which we obtained the data can be found in Kumarathunge et al. 2018, 2019 where the data were originally compiled from the literature.**

specific comments by line number

I recommend rewording the title to clarify what the clause "that affects leaf carbon balance models" applies to:

"A derivation error that affects leaf carbon balance models exists in the current implementation of the Johnson et al. (1942) modified Arrhenius function"

**We thank the reviewer for the comment and have changed the title.**

50: at temperature T

**Done! Line 55**

97, 99: I don't think it's necessary to include the intermediate steps with terms like "DeltaS+-DeltaH" and "298.15—DeltaH". It's also potentially ambiguous to some readers (does "+-" mean "plus or minus", or are the authors including an extraneous plus symbol?)

**We have eliminated the +/- notation, however we retain the intermediate steps for clarity. Line 101 - 117**

The equations, in general: First, the exponential functions with complicated operands should be written out as exp(...) for clarity. Second, the authors should avoid using the "inline" form for the Microsoft Equation Editor equations. The "professional" form is far easier to read. These two issues are particularly bad, for example, in Equation 5.

**We have since reworked the notation so that the equations are more readable. Equations 1 – 10**

115: what does this mean: Data from ... (2019) available from ... (2018)?

**The data are in the 2019 paper, but can be downloaded from a 2018 repository. We have clarified this. Lines 139 - 140**

115: data for what variables?

**Data for A-Ci curves to determine V_cmax_ and J_max_. We have since clarified this. Lines 139 – 140**

125-131: you need to provide a little more info here about the experiment, besides just the species name. What were the treatments and control? what was the context?

**We now provide more information on the background of the experiment. Lines 157 - 166**

138-9: Km is the Michaelis constant for carboxylation specifically (actually the effective Michaelis constant, accounting for competition with O2)

**We have clarified this. Table 1**

In Table 2, the first equation that appears on page 8 appears to be incorrect. I'm not sure if it's a typo. First, even if J = Jmax (in saturating light), Wj should be proportional to the quantity involving ci. Second, even if this was a typo and the closing parenthesis for the min() function was meant to be located after Qin (so that J = min(Jmax, alpha*phi*Qin)), this is a pretty non-standard way to model J, that is, as a simple minimum of capacity and light limited values as opposed to using a non-rectangular hyperbola or something similar. The authors should clarify this equation and its justification.

**We apologize for the error – we have since changed the equations to use the non-rectangular hyperbola model. Table 2**

194-... I don't think the headline result here should be changes in the fitting of each parameter. Since these temperature response equations are invariably applied in a strictly empirical fashion, all that matters is the fit of the overall function.

**We now start with the overall fits of the curves based on BIC and a few example curves. Since equation 3 is categorically wrong due to a derivation error, this demonstrates that equation 10 is preferable. Fig. 2, Lines 212 - 216**

194-... Perhaps the first results graphs should be the actual data for each fitted variable vs temperature, overlaid with the fitted versions of Equations 3 and 10. Yet these relationships don't seem to appear anywhere. That's quite odd.

**We now present the temperature response fits in Figure 2.**

195: which was the y-variable and which the x for these slopes?

**We now make this clear throughout and in the graphs. Lines 2218 – 219, 234, 239, 245, 247, Figs 2 – 4**

313: but nobody does this, arbitrarily holding one parameter constant when fitting a temperature response curve. Unclear why these comparisons are relevant.

**A lot of previous research has done this (i.e. fixing H_d_), in particular the ones addressing thermal acclimation of photosynthetic capacity (e.g. Farquhar et al., 1980, Dreyer et al. 2001, Medlyn et al. 2002, Hikosaka et al. 2006, Kattge & Knorr 2007, Varhammar et al. 2015, Kumarathunge et al. 2019). However, we agree that the fixed H_d_ comparisons are irrelevant (especially since carbon balance models are very sensitive to H_d_, Stinziano et al. 2018) and have dropped that analysis from the manuscript.**

**Dreyer E, Le Roux X, Montpied P, Daudet FA, Masson F. 2001. Temperature response of leaf photosynthetic capacity in seedlings from seven temperate tree species. Tree Physiology 21: 223–232.**

**Farquhar GD, et al. 1980. A biochemical model of photosynthetic CO2 assimilation in leaves of C3 species. Planta 149: 78-90.**

**Hikosaka K, Ishikawa K, Borjigidai A, Muller O, Onoda Y. 2006. Temperature acclimation of photosynthesis: mechanisms involved in the changes in temperature dependence of photosynthetic rate. Journal of Experimental Botany 57: 291–302.**

**Kattge J, Knorr W. 2007. Temperature acclimation in a biochemical model of photosynthesis: a reanalysis of data from 36 species. Plant, Cell & Environment 30: 1176–1190.**

**Kumarathunge DP, Medlyn BE, Drake JE, Tjoelker MG, et al. 2019. Acclimation and adaptation components of the temperature dependence of plant photosynthesis at the global scale. New Phytologist 222:768-784.**

**Medlyn BE, Dreyer E, Ellsworth D, Forstreuter M, Harley PC, Kirschbaum MUF, Le Roux X, et al. 2002. Temperature responses of parameters of a biochemically based model of photosynthesis. II. A review of experimental data. Plant, Cell & Environment 25:1167-1179.**

**Stinziano JR, Way DA, Bauerle WL. 2018. Improving models of photosynthetic thermal acclimation: which parameters are most important and how many should be modified? Global Change Biology 24:1580-1598.

Varhammar A, Wallin G, McLean CM, Dusenge ME, Medlyn BE, Hasper TB, Nsabimana D, Uddling J. 2015. Photosynthetic temperature responses of tree species in Rwanda: evidence of pronounced negative effects of high temperature in montane rainforest climax species. New Phytologist 206: 1000–1012.**

Reviewer: 2

*Comments to the Author*
Summary

The manuscript "A derivation exists in the current implementation of the Johnson et al. (1942) modified Arrhenius function that affects leaf carbon balance models" by Murphy and Stinziano details a study in which the authors find an error in the derivations of the peaked Arrhenius temperature response curve originally presented in Johnson et al. (1942) and simulate its impact on model fitting as well as plant carbon balance models. The authors present the correct derivation and show that the use of the incorrect derivation can lead to errors in model fitting and carbon balance simulations. There is very little to critique about this manuscript. I have checked the derivations and they are sound. The authors have found an error that is important for future studies that use the peaked Arrhenius formula for simulating the temperature response of photosynthetic capacity.

General comments

- The calculation of percent error is not entirely clear. It would be nice to have a more detailed explanation for how the error values were calculated in the statistical analysis portion of the methods.

**We calculated percent error as (slope – 1) * 100%. We now clarify this in the methods. Line 205**

- The point made in the last paragraph of the abstract (L 31-32) was only briefly mentioned in the discussion of the manuscript. It is clear from the study that the revised derivation of the modified Arrhenius model should be used. However, it was not entirely clear why the paradigm should not be used at all (or why the MMRT should be preferred). I think the authors have room to further argue this point, but it seems somewhat tangential to the main goal of the study. As it's currently written, this section distracts more than it adds.

**We have since dropped the discussion of MMRT.**

Minor comments

L23: I would clarify that this is photosynthetic "capacity"

**Done. Line 23**

L37: Removing from what and releasing to what?

**Removing and releasing to the atmosphere – now clarified in text. Lines 38 – 40**

L38: This statement could use a citation.

**Done – added Lombardozzi et al. 2015. Line 41**

L47: I would suggest adding "or equivalently" here.

**Done! Line 51**

LL61-64: It's unclear what you are trying to say here. Are you saying that equation 3 is used for ACi curve fitting? I would suggest rewording this sentence.

**We have reworded this to make it clear that the temperature response function is used to model the temperature response of photosynthetic capacity. Lines 66 - 68**

L67: I would change "state of the Earth system" to "carbon cycling" to be more precise.

**Done. Line 72**

L73: "in calculations covering" is odd phrasing

**We have now rephrased this for clarity. Line –74 - 76**

L93-101: While not entirely necessary, it might be nice to briefly state what is being done in each step here.

**We now provide a brief description for each step. Lines 103 - 119**

L109-111: This is hard to follow. Could you just explain these in complete sentences?

**We have switched to sentences. Lines 131 - 136**

Figure 1: Note what ABQ and LDN are in the legend.

**We now provide a description. Fig. 1**

L186-190: Given what is presented in the Results, there is a lot missing here. Please expand upon what regressions were done and why as well as how the percent changes were calculated.

**We have now provided a substantially larger description of the analyses. Lines –199 - 208**

L326-327: As noted above, it's not entirely clear why MMRT is recommended. What is different? Are parameterizations available? How do the complexities of the two approaches differ? There is a lot to expand upon here and some further expansion is needed if you will want people to follow this recommendation. Alternatively, I think this could be removed and not have much effect on the overall manuscript.

**We have removed all discussion of MMRT from the manuscript.**

Nick Smith, Texas Tech University,

Reviewer: 3

Comments to the Author

The study “A derivation error exists in the current implementation of the Johnson et al. (1942) modified Arrhenius function that affects leaf carbon balance models” by Murphy & Stinziano rederives a version of modified Arrhenius function (as shown in Medlyn et al., 2002) from the original Johnson et al., (1942) functional form.  Authors identified a derivational error between the original Johnson et al., (1942) and Medlyn et al., (2002) functional forms and they found that, this error has impacted the predictions of whole plant carbon balance by causing a deviation of 1.8% between the original and corrected versions of the modified Arrhenius function. However, authors found no significant difference in fitted parameters between the two versions (i.e. original vs corrected model).

I have a few observations the authors may wish to consider.
1. My main frustration is the complete lack of reference to all the other forms of that Johnson et al., (1942) equation available in literature. A number of alternative functions have been used to model the temperature dependences of photosynthetic capacities; Vcmax and Jmax (see, for example, Harley et al. 1986; Long 1991; Harley et al. 1992; Harley & Baldocchi 1995; Lloyd et al. 1995). However, all these equations are actually just alternative expressions of basic Johnson et al., (1942). There is fundamentally no difference between these functions, and it is not clear the real reason for choosing Medlyn et al., (2002) for this study over the others. I would suggest that the authors should also review all the other forms of that equation and verify how different they all are and what fraction of a percent difference in carbon balance they make. I firmly think that such an evaluation is inevitable.

**We thank the reviewer for the references and comments. We chose the Medlyn et al. 2002 form because that is the form that is used in models and acclimation studies (e.g. Kattge & Knorr, 2007; Kumarathunge et al., 2019). We now briefly review the other forms of the equation. We note that: 1) with minimal algebra, Harley & Baldocchi 1995 and Lloyd 1995 are identical to the Medlyn et al. 2002 equations, and 2) Long 1991 does not actually provide a peaked form of the Arrhenius equation. We now show the derivations of Harley et al. 1992 Farquhar et al. 1980 to show that they are identical to Medlyn et al 2002 when relativized to temperature in Appendix A. We also note that Harley et al. 1986, relativizes to the same structure as our equation in Appendix A. Therefore, we simply need to compare Equations 3 and 10 from our manuscript. Lines 123 – 131, Appendix A**

2. In the process of rederivation of the modified Arrhenius function, the authors defined that the term c’’ of the Johnson et al., (1947) as a second derivative of the rate and assumed it as a temperature independent constant (L89). If the rate of a given reaction is temperature dependent, it is not clear how the second derivative is not temperature dependent given the term c’’ is not explicitly defined in Johnson et al., (1942). Hence, it is not clear how the term c’’ cancelled out when relativizing the function to a reference temperature (compare Equations 6 & 7).

**We revisited the original paper, and it is clear from our re-reading that c’’ is actually a constant, not a second derivative. We have since changed the text. Line 94**

3. When comparing the fitted temperature response parameters between the corrected and Medlyn et al., (2002) derivation of the modified Arrhenius function, authors found that most parameters are not significantly different, though small errors still exist. If the parameters are not different between two versions of the function, can those existing small errors due to a random chance? I would like to see how the fitted temperature response curves differ between the two functional forms, perhaps a figure that shows a sample dataset of Vcmax or Jmax and fitted curves for both functions overlaid on the same panel. If anything, this analysis will tell us that both functions perform equally well (or not). This may repeat with other versions of the Johnson et al., (1942) (see my comment 1).  Further, I would like to see how those “small errors” due to the missing term in Medlyn et al., (2002) scaled at stand or regional scale GPP estimates of a stand scale model or a terrestrial biosphere model. Currently, it is not convincing how such a small (1.8%) deviation in daily carbon balance really matter in modelling GPP at larger scales.

**We took a more robust approach to curve fitting during the revisions. Originally, we were only able to fit 92 paired V_cmax_/J_max_ curves – this was because we used a single set of starting parameters. This time, we use 1,000 sets of starting parameters, expanding the successful curve fits to 196 complete pairs (~1/2 of candidate temperature responses) (Lines 144 – 153). In the new analysis, E_a_ shows a large difference between the two equations, with the other parameters essentially identical (lines 221 – 224, Fig. 3). We now also show examples of temperature responses where the curves are nearly identical, and examples where they are different (Fig. 2). Our modeling now shows ~14-16% difference in *daily* net plant carbon gain due to the expanded and more robust set of curve fits (lines 232 – 248, Fig. 4). The point of this manuscript was to show the impact on a whole plant level, not at the stand level. However, if we assume 1.8% difference in daily carbon balance, and use a compound interest equation, P = A (1 + r) ^ t, where P is the endpoint value, A is the original amount (assume 1 g in this case for simplicity), r is the growth rate (1.8% day^-1^ in this case), and t is the time period over which interest is applied (days in this case), then after 30 days, P increases from 1 to 1.71, indicating a 71% difference after one month for a single plant. Therefore, even the original 1.8% difference in *daily* carbon balance is very large when compounded day after day. However, after the revisions, the difference is nearly 10-fold larger and we believe that the results at the whole plant level are sufficient to make our point that the equation needs to be corrected.**

4. L329-335 - I appreciate the arguments that authors have made here. But this manuscript not directly discuss or justify reasons for moving beyond the modified Arrhenius paradigm for modelling temperature response of process rates. Despite the so called “missing term” as identified by authors (but see my point no 2), the modified Arrhenius function and its alternative expressions have long been used as a robust function to quantify temperature response of photosynthetic capacities (Dreyer et al., 2001; Medlyn et al., 2002; Hikosaka et al., 2006; Kattge & Knorr 2007; Varhammar et al., 2015; Kumarathunge et al., 2019).  I believe that empirical approaches based on the original Johnson et al., (1942); such as Kattge & Knorr (2007) and updated recently by Kumarathunge et al (2019) do provide a robust formulation to quantify photosynthetic capacities across the full biological range of temperatures.

**In accordance with Reviewer 2’s comments, we have eliminated the discussion of moving beyond the modified Arrhenius paradigm. While we agree that the modified Arrhenius equation appears robust, its current form is categorically wrong and in error. The empirical approaches “based on the original Johnson et al” equation are technically not based on that equation because a term was erroneously dropped. Presumably by rectifying this error, the equation will prove even more robust in the future.**

Minor 
L 91 – Correct units of ΔS: J mol-1 K-1 (or kJ mol-1 K-1)

**Done! Line 95**

L132-134 - Is there evidence for a connection between the ratio of photosynthesis to respiration alter the temperature response of photosynthetic capacities (or carbon balance)?

**We have not specifically investigated this. Rather, we wanted to demonstrate that even though the impact of the equation switch impacted just the photosynthesis part of our modeling, there was a substantial impact on *net carbon balance* under both scenarios of low and high respiration, where the impact on photosynthesis may be washed out in the net calculations.**
