## Supplementary material for "A derivation error that affects carbon balance models exists in the current implementation of the modified Arrhenius function": R package for analysis: Murphy_Stinziano_SI_Appendix.docx (from another version of this preprint: version 2, 2020-04-23)

**Running Title: Derivation error in modified Arrhenius model**

Author: Bridget Murphy^1^, Joseph R. Stinziano^2,*^

Affiliations: ^1^ Department of Biology, University of Western Ontario, London, ON, Canada;
^2^ Department of Biology, University of New Mexico, Albuquerque, NM, USA

**Supplementary information**

**Appendix A – Alternate derivations of the modified Arrhenius equation**

Below are the temperature relativizations of Farquhar et al. (1980), Harley et al. (1986), and Harley et al. (1992) showing that Farquhar et al. (1980) and Harley et al. (1992) can be transformed to be identical to Equation 3 in the manuscript, while Harley et al. (1986) can be transformed to be identical to Equation 10 in the manuscript.

*Farquhar et al. (1980)*

From Eq. 36 in Farquhar et al. (1980):

$k=\frac{c\times exp[-E/(RT_{k})]}{1+exp[(ST_{k}-H)/(RT_{k})]}$	Equation A1

Relativizing to 25 °C:

$k/k_{25}=\left[ \frac{c\times exp[-E/(RT_{k})]}{1+exp[(ST_{k}-H)/(298.15R)]} \right]\left[ \frac{1+exp[(ST_{k}-H)/(298.15R)]}{c\times exp[-E/(T_{k}R)]} \right]$	Equation A2

Simplifying:

$k/k_{25}=exp[E/(298.15R)-E/(RT_{k})]\left[ \frac{1+exp[(S298.15-H)/(298.15R)]}{1+exp[(ST_{k}-H)/(RT_{k})]} \right]$	Equation A3

$k=k_{25}\exp\left[ E\frac{\left( T_{k}-298.15 \right)}{\left( 298.15RT_{k} \right)} \right]\left[ \frac{1+exp[(298.15S-H)/(298.15R)]}{1+exp[(ST_{k}-H)/(RT_{k})]} \right]$	Equation A4

where E is activation energy (E_a_), S is deactivation entropy (ΔS), and H is deactivation energy (H_d_).

*Harley et al. (1986)*

From Eq. 7 in Harley et al. (1986):

$k=\frac{T_{k}exp[c-{\Delta H}_{a}/(RT_{k})]}{1+exp[(\Delta ST_{k}-{\Delta H}_{d})/(RT_{k})]}$	Equation A5

Relativizing to 25 °C:

$k/k_{25}=\left[ \frac{T_{k}exp[c-{\Delta H}_{a}/(RT_{k})]}{1+exp[(\Delta ST_{k}-{\Delta H}_{d})/(RT_{k})]} \right]\left[ \frac{1+exp[(\Delta S298.15-{\Delta H}_{d})/(R298.15)]}{298.15exp[c-{\Delta H}_{a}/(R298.15)]} \right]$	Equation A6

Simplifying:

$k/k_{25}=\frac{T_{k}}{298.15}exp[{\Delta H}_{a}/(R298.15)-{\Delta H}_{a}/(RT_{k})]\left[ \frac{1+exp[(\Delta S298.15-{\Delta H}_{d})/(R298.15)]}{1+exp[(\Delta ST_{k}-{\Delta H}_{d})/(RT_{k})]} \right]$	Equation A7

$k=k_{25}\frac{T_{k}}{298.15}\exp\left[ {\Delta H}_{a}\frac{\left( T_{k}-298.15 \right)}{\left( 298.15RT_{k} \right)} \right]\left[ \frac{1+exp[(298.15\Delta S-{\Delta H}_{d})/(298.15R)]}{1+exp[(\Delta ST_{k}-{\Delta H}_{d})/(RT_{k})]} \right]$	Equation A8

where ΔH_a_ is activation energy (E_a_), and ΔH­_d_ is deactivation energy (H_d_).

*Harley et al. (1992)*

From Eq. 9 in Harley et al. (1992):

$k=\frac{exp[c-{\Delta H}_{a}/(RT_{k})]}{1+exp[(\Delta ST_{k}-{\Delta H}_{d})/(RT_{k})]}$	Equation A9

Relativizing to 25 °C:

$k/k_{25}=\left[ \frac{exp[c-{\Delta H}_{a}/(RT_{k})]}{1+exp[(\Delta ST_{k}-{\Delta H}_{d})/(RT_{k})]} \right]\left[ \frac{1+exp[(\Delta S298.15-{\Delta H}_{d})/(R298.15)]}{exp[c-{\Delta H}_{a}/(R298.15)]} \right]$	Equation A10

Simplifying:

$k/k_{25}=exp[{\Delta H}_{a}/(R298.15)-{\Delta H}_{a}/(RT_{k})]\left[ \frac{1+exp[(\Delta S298.15-{\Delta H}_{d})/(R298.15)]}{1+exp[(\Delta ST_{k}-{\Delta H}_{d})/(RT_{k})]} \right]$	Equation A11

$k=k_{25}\exp\left[ {\Delta H}_{a}\frac{\left( T_{k}-298.15 \right)}{\left( 298.15RT_{k} \right)} \right]\left[ \frac{1+exp[(298.15\Delta S-{\Delta H}_{d})/(298.15R)]}{1+exp[(\Delta ST_{k}-{\Delta H}_{d})/(RT_{k})]} \right]$	Equation A12

where ΔH_a_ is activation energy (E_a_), and ΔH­_d_ is deactivation energy (H_d_).

**Supplementary Information**

*Instructions for installing {arrhenius.comparison} in R for review purposes*

1. Download the arrhenius.comparison.tar.gz file
2. Set working directory in R to the directory that holds the arrhenius.comparison.tar.gz file using setwd()
3. Run:
   install.packages("arrhenius.comparison_1.0.1.tar.gz", repos = NULL, type = "source")
   library(arrhenius.comparison)
